# A meta-analysis of *in vitro* release of hydrophilic therapeutics from contact lenses using mathematical modeling

**DOI:** 10.1101/2025.07.11.664435

**Authors:** Lucia Carichino, Kara L. Maki, Narshini D. Gunputh, Chau-Minh Phan

**Affiliations:** School of Mathematics and Statistics, Rochester Institute of Technology, 85 Lomb Memorial Drive, Rochester, 14623, NY, USA; Centre for Ocular Research & Education (CORE), School of Optometry & Vision Science, University of Waterloo, 200 University Avenue West, Waterloo, N2L 3G1, ON, Canada; Centre for Eye and Vision Research (CEVR), Unit 901-903, 17W Hong Kong Science Park, Shatin, Hong Kong

**Keywords:** therapeutic release, contact lens, diffusion coefficient, ophthalmic drug delivery

## Abstract

A meta-analysis was conducted to study the *in vitro* release of hydrophilic therapeutics from contact lenses. Fifty-two experiments were studied that measure the cumulative release of therapeutics from (mostly) commercial contact lenses placed in a vial. A mathematical model and a parameter fitting algorithm is presented to estimate the diffusion coefficient (*D*) and 50% therapeutic release time (*T*_50_) of all the experimental lens-therapeutic combinations. The mathematical framework was validated against previous studies. Statistical methods were used to analyze the relationships between lens materials, therapeutic properties, and predicted parameter values (*D* and *T*_50_). It was found that lens water content directly and moderately influences the estimated diffusion coefficient. More specifically, the median diffusivity of silicone hydrogel (SH) contact lenses was statistically different from conventional hydrogel (CH) lenses. Other lens and therapeutic properties dependencies on diffusivity were complex with special cases studied to elicit dependencies. A predictive tool was constructed to estimate the logarithm of 50% therapeutic release time, log(*T*_50_), given the lens water content and the therapeutic molecular volume and density. The statistical model explained 64% of the variability of the log(*T*_50_) and can be used in the preliminary stages of contact lens drug delivery development.

## 1. Introduction

Vision impairment, commonly caused by glaucoma, diabetic retinopathy, cataracts, and refractive error, affected 1.1 billion people worldwide in 2020, with the number expected to rise to 1.8 billion by 2050 (Swenor and Ehrlich, 2021). Drugs for treating these diseases are usually administered by eye drops (Xu et al., 2018). However, while eye drops are cost-effective and convenient, they suffer from low bioavailability due to spillage, dilution, blinking, and tear drainage after being administered to the eye (Xu et al., 2018; Guzman-Aranguez et al., 2013). One approach that has been proposed to improve drug delivery to the eye is with soft contact lenses (Ciolino et al., 2009). These devices can not only act as a reservoir to provide sustained drug release, but can also effectively shield the drugs from the various aforementioned ocular removal mechanisms (Jones et al., 2021; Fan et al., 2020). Therefore, drug delivery via soft contact lenses can increase drug retention and bioavailability on the ocular surface (Guzman-Aranguez et al., 2013; Jones et al., 2021).

Commercial contact lenses have continuously evolved with the first soft contact lens approved by the FDA in 1971 (Fan et al., 2020; Moreddu et al., 2019). The first soft contact lenses manufactured were conventional hydrogel (CH) contact lenses (Fan et al., 2020; Moreddu et al., 2019). CH contact lenses differ in water content, as water content is important for on-eye performance and oxygen permeability, and ionicity (Fan et al., 2020; Moreddu et al., 2019). Silicone hydrogel (SH) contact lenses, the next class of soft contact lenses launched in 1999, were introduced to increase the oxygen permeability (Fan et al., 2020; Moreddu et al., 2019). Subsequent generations of SH contact lenses aim to decrease the stiffness and increase wettability of the lens caused by the additional silicone (Fan et al., 2020). Commercial contact lenses are safe and easy to manufacture, making them a natural choice for the delivery of ophthalmic drugs (Fan et al., 2020).

Research on the use of contact lenses as a drug delivery vehicle has increased drastically in the past decade (Jones et al., 2021). Acuvue Theravision, produced by Johnson & Johnson Vision Care, Inc., a ketotifen-releasing contact lens, has been approved by the U.S. Food and Drug Administration (FDA) in 2022 (Johnson and Vision). However, issues with contact lens drug delivery still remain, including burst release within the first hours or the inability to maintain sustained therapeutic release over the desired treatment period, warranting further investigation (Jones et al., 2021; Silbert, 1996; Fan et al., 2020; Ciolino et al., 2009).

Designing a contact lens to deliver a specific therapeutic at a specific dosage requires a predictable and controlled release. *In vitro* experiments, evaluating drug delivery from contact lenses using a vial, have shown that several factors can affect the drug release kinetics. SH lenses, which have a lower water content than CH lenses, tended to absorb fewer therapeutics than CH lenses, and consequently release less (Karlgard et al., 2003). Among SH lenses, those containing a higher portion of hydrophilic phases absorbed more therapeutics and released them faster (Xu et al., 2011). For positively charged drugs, such as anti-muscarinic drugs atropine and pirenzepine, it was found that contact lens materials with a surface charge or ionicity released more drugs than nonionic lenses (Hui et al., 2017). Consequently, exploiting electrostatic interactions has been proposed to extend the therapeutic release duration from commercially available SH lenses (Torres-Luna et al., 2020). In addition, the environment in which the lenses were placed also played a role. For example, more sophisticated *in vitro* eye models, with low but constant fluid flow, showed that the release of a therapeutic from a contact lens in that environment is much slower than compared to a vial (Bajgrowicz et al., 2015; Phan et al., 2016, 2021).

Mathematical models have been used to predict the release of a therapeutic from a contact lens and to improve lens delivery. Some models consider *in vitro* (Ali et al., 2007; Dixon and Chauhan, 2017; Lanier et al., 2021; Pimenta et al., 2016; Peng et al., 2010; Peng and Chauhan, 2011) release while other consider *in vivo* release (Li and Chauhan, 2006; Anderson and Luke, 2024; Toffoletto et al., 2023; Gause et al., 2016). The theoretical studies focused on *in vitro* release modeled the release of a loaded therapeutic from a contact lens via diffusion (Torres-Luna et al., 2020; Li and Chauhan, 2006; Peng et al., 2010; Peng and Chauhan, 2011; Pimenta et al., 2016; Lanier et al., 2021). In general, these works estimated the diffusivity of lens-therapeutic combinations, but they did not consider how the predicted diffusivity changes with a wide range of lens and therapeutic properties.

The aim of this study is to conduct a meta-analysis to characterize the dependence of lens and therapeutics properties on the release kinetics of hydrophilic therapeutics from pre-soaked contact lenses placed in a vial of fluid. Fifty-two experiments were analyzed using a mathematical framework, to estimate the diffusion coefficient of the lens-therapeutic combination, and a statistical framework, to understand variations in the estimated diffusion coefficients. An understanding of therapeutic transport can help facilitate the development of better materials for ocular drug delivery.

## 2. Methods

A schematic of the *in vitro* therapeutic delivery analysis is shown in Figure 1. Experimental data, a mathematical model, and a fitting algorithm were used in synergy to estimate the diffusivity and the release time of 50% of a therapeutic from a contact lens in a vial. Predicted diffusivities and release times were then analyzed in terms of lens and therapeutics properties using statistical methods. Section 2.1 introduces the experimental data. Section 2.2 explains the math model used in parameter estimation algorithm described in Section 2.3. Lastly, in Section 2.4, the \statistical methods used to conduct the meta-analysis are introduced.

**Figure 1:**
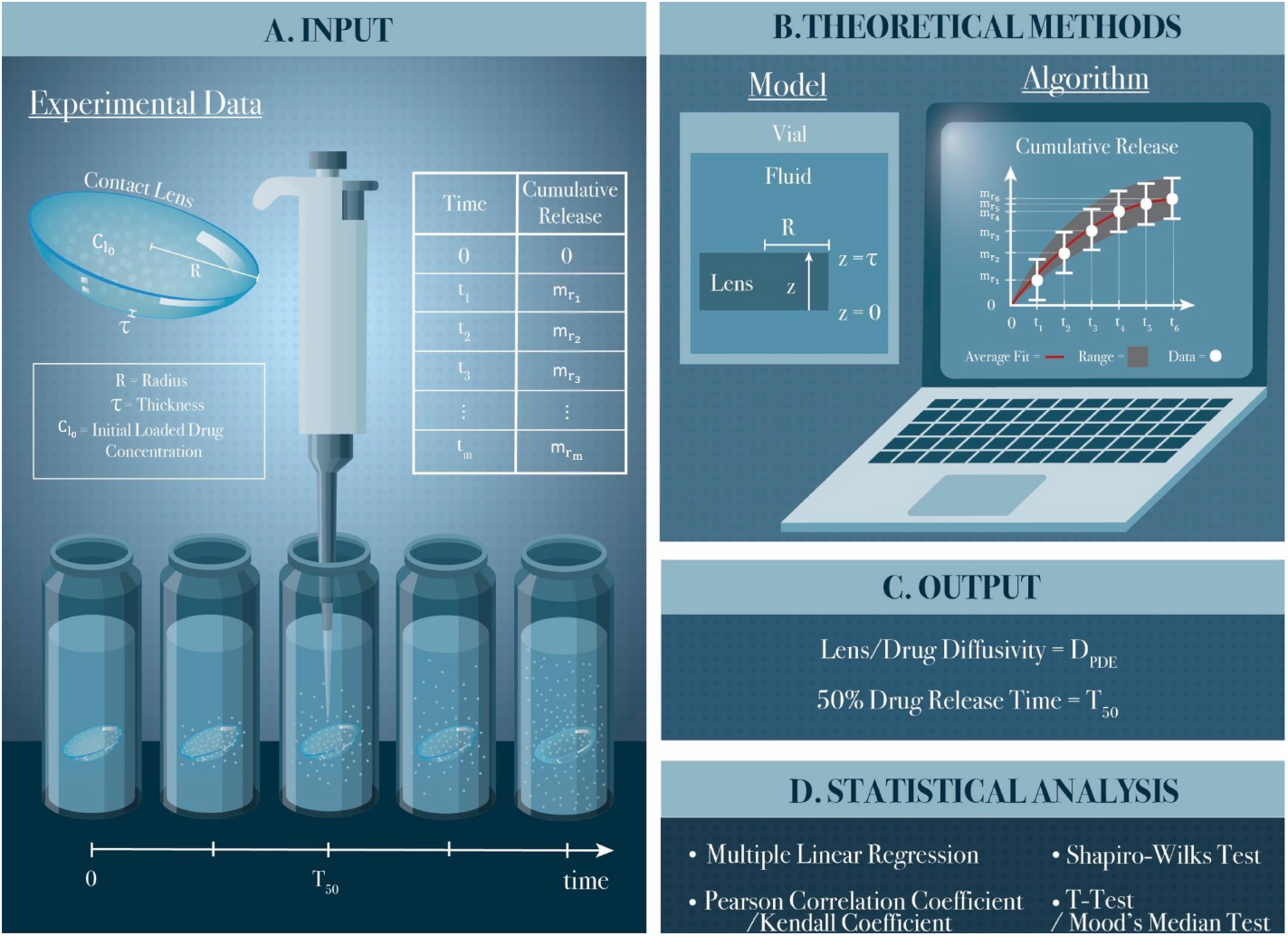
Schematic of the *in vitro* therapeutic delivery analysis performed. Starting from (**A**) experimental data, (**B**) a mathematical model and a fitting algorithm were used to estimate (**C**) the diffusivity and the release time of 50% of a therapeutic from a contact lens in a vial. (**D**) Predicted diffusivities and 50% release times were then analyzed using statistical methods. Illustration created by Kerstyn Gay (Gay).

### 2.1. Experimental Data

The cumulative therapeutic release time series datasets for this study were taken from eleven different publications listed in chronological order in Table 1. *In vitro* experiments considered measured the release of hydrophilic therapeutics from (mostly) commercial contact lenses placed in a vial. The therapeutic was loaded into the contact lens by soaking method. Such experiments most closely satisfied the simplifying assumptions used in the still-to-be-introduced mathematical model. In what follows, the general experiment procedure, and the experimental data used for this study, are explained in Section 2.1.1. Then, in Section 2.1.2, the partition coefficient estimates are presented. Finally, the different lens and therapeutic properties are characterized in Section 2.1.3.

**Table 1:**
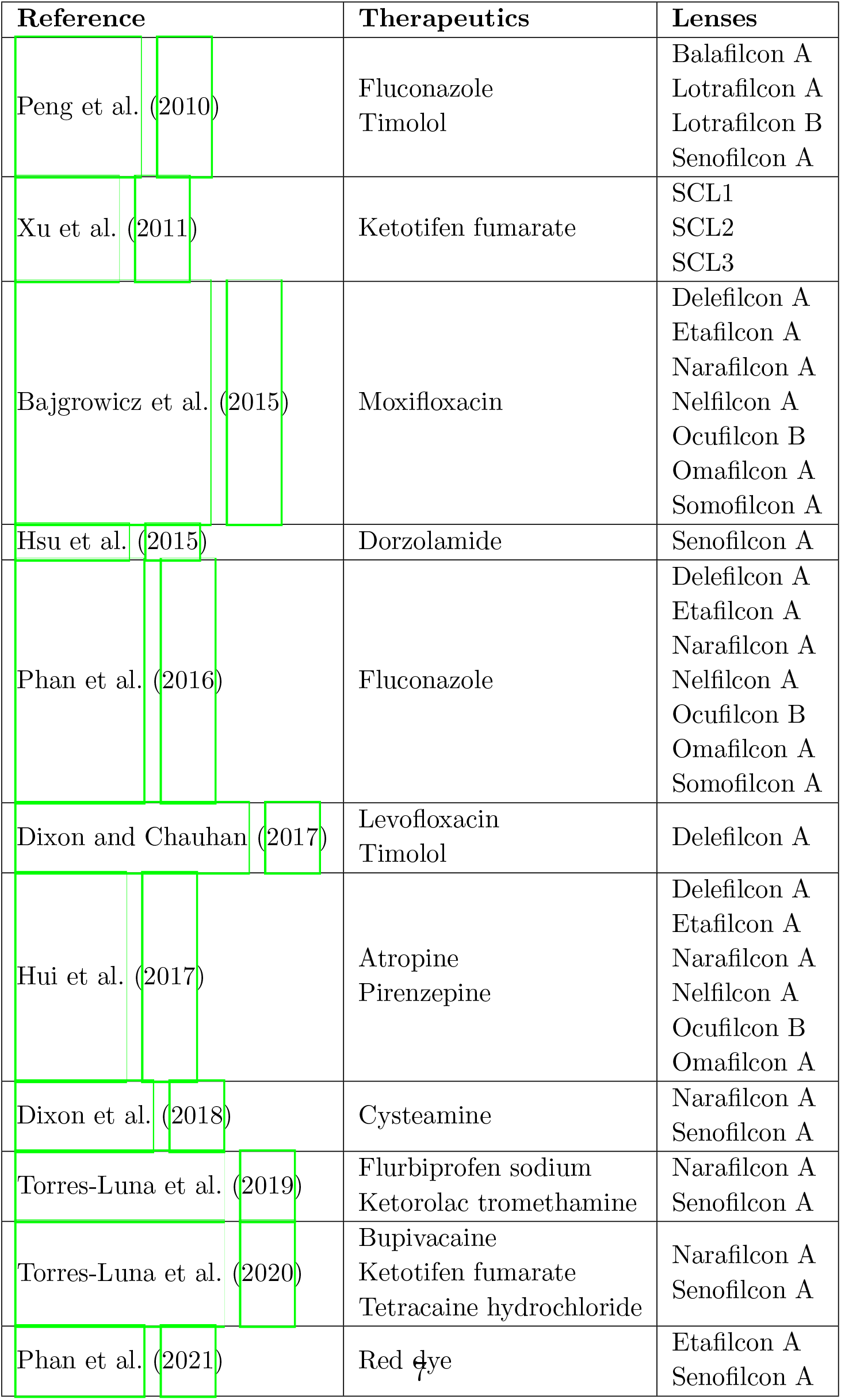
List of publications from which the datasets have been extracted, with details on the contact lenses and therapeutics considered. For the first publication, the contact lens balaficon A is only loaded with timolol.

| Reference | Therapeutics | Lenses |
| --- | --- | --- |
| Peng et al. (2010) | Fluconazole<br>Timolol | Balafilcon A<br>Lotrafilcon A<br>Lotrafilcon B<br>Senofilcon A |
| Xu et al. (2011) | Ketotifen fumarate | SCL1<br>SCL2<br>SCL3 |
| Bajgrowicz et al. (2015) | Moxifloxacin | Delefilcon A<br>Etafilcon A<br>Narafilcon A<br>Nelfilcon A<br>Ocufilecon B<br>Omafilcon A<br>Somofilcon A |
| Hsu et al. (2015) | Dorzolamide | Senofilcon A |
| Phan et al. (2016) | Fluconazole | Delefilcon A<br>Etafilcon A<br>Narafilcon A<br>Nelfilcon A<br>Ocufilecon B<br>Omafilcon A<br>Somofilcon A |
| Dixon and Chauhan (2017) | Levofloxacin<br>Timolol | Delefilcon A |
| Hui et al. (2017) | Atropine<br>Pirenzepine | Delefilcon A<br>Etafilcon A<br>Narafilcon A<br>Nelfilcon A<br>Ocufilecon B<br>Omafilcon A |
| Dixon et al. (2018) | Cysteamine | Narafilcon A<br>Senofilcon A |
| Torres-Luna et al. (2019) | Flurbiprofen sodium<br>Ketorolac tromethamine | Narafilcon A<br>Senofilcon A |
| Torres-Luna et al. (2020) | Bupivacaine<br>Ketotifen fumarate<br>Tetracaine hydrochloride | Narafilcon A<br>Senofilcon A |
| Phan et al. (2021) | Red dye | Etafilcon A<br>Senofilcon A |

#### 2.1.1. Cumulative Therapeutic Release Experiment

Studies, referenced in Table 1, measured (in mass) the cumulative release of different therapeutic from SH and CH contact lenses after being placed in a vial of fluid (see Figure 1**A**). The therapeutic *in vitro* experiment had two parts: soaking and release in vials. First, a therapeutic was loaded into a contact lens by the *soaking* method. To do so, a contact lens was placed in a vial containing a reported therapeutic concentration, called the loading therapeutic concentration, denoted by 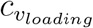, and reported in Appendix A Table A.8, for a specified amount of time. Then, the therapeutic loaded contact lens was placed in a vial of phosphate buffered saline (PBS) with volume *v*_*v*_, reported in Appendix A Table A.8. The amount of therapeutic *released* from the contact lens into the vial was measured over a twenty-four hour period. This is the measured culminate therapeutic release.

For each lens-therapeutic pair, the experiment was run multiple times; consequently, the cumulative therapeutic release time series datasets were reported as sample means with associated standard derivations. The experimental time series were denoted by 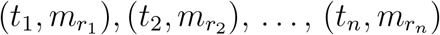, where *t*_*i*_ is the time of the *i*-th measurement, 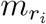 is either the measured sample mean therapeutic mass in the vial or the measured sample mean mass plus or minus its sample standard derivation, and *n* is the total number of measurements. An example cumulative therapeutic release time series dataset is shown in Figure 1**B**, where the circles denotes sample mean and the bars indicate plus or minus one standard derivation. Experimental datasets were either shared directly by the authors or were extracted from the publication.

#### 2.1.2. Estimated Partition Coefficient

The partition coefficient, *K*, is defined to be the ratio of the lens therapeutic concentration to the vial therapeutic concentration at equilibrium (end of soaking). The partition coefficient can be estimated by measuring the initial and final vial therapeutic concentrations during soaking (Dixon and Chauhan, 2017). However, Dixon and Chauhan (2017) found for hydrophilic therapeutics changes in the therapeutic concentration of the soaking solution could not be measured experimentally. Consequently, for experiments releasing hydrophilic therapeutics, Dixon and Chauhan (2017) characterized the partition coefficient as follows:

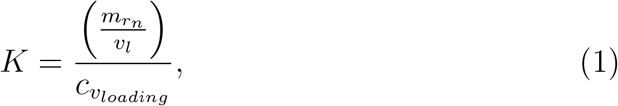

where 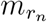 is final measured therapeutic mass in vial released from the contact lens, v_l_ is an estimated lens volume, and 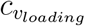 is the loading therapeutic concentration. That is, the lens therapeutic concentration at the end of soaking was estimated by the lens therapeutic concentration released. Consequently, it was assumed that all the drug loaded into the contact lens was released in the subsequent experiment. Table A.8 in Appendix A reports the estimated partition coefficients for all lens-therapeutic combinations considered in this work using Eq. (1).

#### 2.1.3. Lens and Therapeutic Properties

A total of fourteen contact lenses were investigated in combination with different therapeutics, as described in Table 1. The key lens properties considered were lens material (SH vs CH), the FDA Group (Groups I-V) (Food & Drug Administration, FDA; Green, 2018; Rhea, 2017), the ionicity (Food & Drug Administration, FDA; Green, 2018; Rhea, 2017), the water content, the thickness, *τ*, and the radius, *R*. FDA Groups were defined as follows: (I) non-ionic, low water content (< 50%) CH lenses; (II) non-ionic, high water content (> 50%) CH lenses; (III) ionic, low water content CH lenses; (IV) ionic, high water content CH lenses; and (V) SH lenses (Hutter et al., 2012). In the datasets considered, SH lenses were classified as Group V, and CH lenses into Groups II and IV, so no lenses from Group I and III were considered. A summary of the fourteen contact lens properties, organized by FDA Group, is provided in Table 2.

**Table 2:** Description of contact lenses studied and their properties. Abbreviations are as follows: EGDMA: Ethylene glycol dimethacrylate, FMA: N-Formylmethyl acrylamide, PVP: Polyvinyl pyrrolidone, NVP: N-vinyl pyrrolidone, DMA: N,N-Dimethylacrylamide, MPDMS: monofunctional polydimethylsiloxane, TEGDMA: Tetraethleneglycol dimethacrylate. Symbols indicate references: *a* = Bajgrowicz et al. (2015), *b* = Phan et al. (2021), *c* = Chou (2008), *d* = Peng et al. (2010), *e* = Chatterjee et al. (2020), and *f* = Xu et al. (2011).

| Contact lens<br>Monomers | Commercial<br>name | FDA<br>group | Ionic | Water<br>content<br>[%] | Thickness<br>$\tau$<br>[mm] | Radius<br>$R$<br>[mm] |
| --- | --- | --- | --- | --- | --- | --- |
| Nelfilcon A<br>PVA, FMA,<br>PEG<sup>a</sup> | DAILIES<br>Aquacomfort<br>Plus | II | No | 69<sup>a</sup> | 0.100<sup>a</sup> | 7.000<sup>a</sup> |
| Omafilcon A<br>HEMA, PC,<br>EGDMA<sup>a</sup> | Proclear<br>1-Day | II | No | 62<sup>a</sup> | 0.090<sup>a</sup> | 7.100<sup>a</sup> |
| Etafilcon A<br>HEMA, MAA<sup>a</sup> | 1-Day<br>Acuvue Moist | IV | Yes | 58<sup>a</sup> | 0.080<sup>a,b</sup> | 7.150<sup>a,b</sup> |
| Ocufilcon B<br>HEMA, PVP,<br>MAA<sup>a</sup> | Biomedics<br>1-Day | IV | Yes | 53<sup>a</sup> | 0.070<sup>a</sup> | 7.100<sup>a</sup> |
| Balafilcon A<br>TRIS, NVP<sup>c</sup> | Pure<br>Vision 2 | V | Yes | 36<sup>d</sup> | 0.090<sup>d</sup> | 6.250<sup>d</sup> |
| Delefilcon A<br>Not<br>disclosed<sup>a</sup> | DAILIES<br>TOTAL 1 | V | No | 33<sup>a</sup> | 0.090<sup>a</sup> | 7.050<sup>a</sup> |
| Lotrafilcon A<br>DMA, TRIS<br>Siloxane<sup>e</sup> | NIGHT<br>& DAY | V | No | 24<sup>d</sup> | 0.080<sup>d</sup> | 6.390<sup>d</sup> |
| Lotrafilcon B<br>DMA, TRIS<br>Siloxane<sup>e</sup> | O<sub>2</sub> Optix | V | No | 33<sup>d</sup> | 0.080<sup>d</sup> | 6.390<sup>d</sup> |
| Narafilcon A<br>HEMA, DMA,<br>TEG, Siloxane,<br>MPDMS, PVP<sup>a</sup> | 1-Day<br>ACUVUE<br>TruEye | V | No | 46<sup>a</sup> | 0.090<sup>a</sup> | 7.100<sup>a</sup> |
| SCL1<br>DMA, TRIS,<br>EGDMA,<br>MA-PDMS-MA<sup>f</sup> | - | V | - | 38<sup>f</sup> | 0.084<sup>f</sup> | 6.005<sup>f</sup> |
| SCL2<br>DMA, TRIS,<br>EGDMA,<br>MA-PDMS-MA<sup>f</sup> | - | V | - | 30<sup>f</sup> | 0.082<sup>f</sup> | 5.970<sup>f</sup> |
| SCL3<br>DMA, TRIS,<br>EGDMA,<br>MA-PDMS-MA<sup>f</sup> | - | V | - | 25<sup>f</sup> | 0.081<sup>f</sup> | 5.880<sup>f</sup> |
| Senofilcon A<br>HEMA, DMA,<br>Siloxane,<br>TEGDMA,<br>PVP, MPDMS<sup>a</sup> | Acuvue Oasys | V | No | 20<sup>a,d</sup> | 0.084<sup>a,d</sup> | 7.150<sup>a,d</sup> |
| Somofilcon A<br>Not disclosed<sup>a</sup> | Clariti 1-Day | V | - | 56<sup>a</sup> | 0.070<sup>a</sup> | 7.050<sup>a</sup> |

The properties of the therapeutics examined are reported in Table 3, which include molecular mass, density, molecular volume, and charge. The reported molecular mass and density were found in the literature, and molecular volume was computed by dividing the molecular mass by the density.

**Table 3:** Description of therapeutics used and their respective properties. Symbols denote references: *a* = PubChem, *b* = ChemSpider, *c* = Thal et al. (2016), *d* = Lui et al. (1998), *e* = Torres-Luna et al. (2020), *f* = El-Housiny et al. (2018), *g* = Sciences, *h* = Uivarosi (2013), *i* = Chavez-Bravo et al. (2016), *j* = ChemicalBook, *k* = Kim et al. (2018), *l* = Kim and Na (2019), *m* = Torres-Luna et al. (2019), *n* = Shahid et al. (2022), *o* = Loftsson (2014), *p* = Tatarchuk et al. (2023), *q* = ChemSrc, and *r* = Phan et al. (2021).

| Therapeutic | Molecular mass<br>[g/mol] | Density<br>[g/cm<sup>3</sup>] | Molecular volume<br>[cm<sup>3</sup>/mol] | Charge |
| --- | --- | --- | --- | --- |
| Atropine | 289.37<sup>a</sup> | 1.20<sup>b</sup> | 241.14 | Positive<sup>c</sup> |
| Bupivacaine | 342.90<sup>a</sup> | 0.99<sup>d</sup> | 343.07 | Positive<sup>e</sup> |
| Cysteamine | 77.15<sup>a</sup> | 1.00<sup>b</sup> | 77.15 | Positive<sup>l</sup> |
| Dorzolamide | 324.44<sup>a</sup> | 1.6<sup>b</sup> | 202.78 | Positive<sup>o</sup> |
| Fluconazole | 306.27<sup>a</sup> | 1.50<sup>b</sup> | 204.18 | Negative<sup>f</sup> |
| Flurbiprofen sodium | 244.26<sup>a</sup> | 1.01<sup>b</sup> | 240.42 | Negative<sup>c</sup> |
| Ketotifen fumarate | 309.43<sup>a</sup> | 0.97<sup>g</sup> | 319.66 | Positive<sup>e</sup> |
| Ketorolac tromethamine | 255.27<sup>a</sup> | 1.33<sup>q</sup> | 191.93 | Positive<sup>m,n</sup> |
| Levofloxacin | 361.37<sup>a</sup> | 1.5<sup>b</sup> | 240.19 | Positive<sup>p</sup> |
| Moxifloxacin | 401.43<sup>a</sup> | 1.40<sup>b</sup> | 286.74 | Positive<sup>h</sup> |
| Pirenzepine | 351.40<sup>a</sup> | 1.30<sup>b</sup> | 270.31 | Positive<sup>c</sup> |
| Red dye | 650.60<sup>a</sup> | 1.00<sup>t</sup> | 650.60 | Negative<sup>i</sup> |
| Tetracaine hydrochloride | 200.80<sup>a</sup> | 1.13<sup>j</sup> | 178.00 | Positive<sup>e</sup> |
| Timolol | 432.50<sup>a</sup> | 1.20<sup>b</sup> | 360.40 | Positive<sup>k</sup> |

### 2.2. Mathematical Model

A mathematical model of therapeutics release from a contact lens was created. Two therapeutics concentration were considered: the lens therapeutic concentration, denoted by *c*_*l*_, and the vial therapeutic concentration, denoted by *c*_*v*_. The lens therapeutic concentration variations were neglected in the radial direction; i.e., *c*_*l*_(*z, t*) was assumed to vary only along the thickness of the lens, represented by the variable *z* (see Figure 1**B**) and time *t*. The contact lens was assumed to be loaded with a dilute, spatially independent concentration of therapeutic, 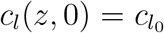, and placed into a vial of volume *v*_*v*_ containing no therapeutic. That is, the vial therapeutic concentration *c*_*v*_ was zero at t = 0; i.e., *c*_*v*_| _*t*=0_ = 0.

At the lens surface, the lens therapeutic concentration was assumed to be proportional to the vial therapeutic concentration, *c*_*l*_ = *Kc*_*v*_, where *K* is the partition coefficient defined in Eq. (1) (Dixon and Chauhan, 2017; Lanier et al., 2021). Because vial volume, *v*_*v*_, was much larger than the lens volume, *v*_*l*_, was assumed *c*_*v*_ ≈ 0 as argued in Appendix B. Therefore, sink boundary conditions were assumed. That is,

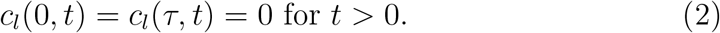

The assumption of sink boundary conditions was also used in prior works (Siepmann and Siepmann, 2012; Peng and Chauhan, 2011; Dixon and Chauhan, 2017; Dixon et al., 2018).

The transport of the lens therapeutic was assumed to be governed by Fick’s law of diffusion (Li and Chauhan, 2006; Peng et al., 2010; Peng and Chauhan, 2011; Pimenta et al., 2016; Lanier et al., 2021). Consequently, lens therapeutic concentration, *c*_*l*_*(z, t)*, was assumed to be governed by

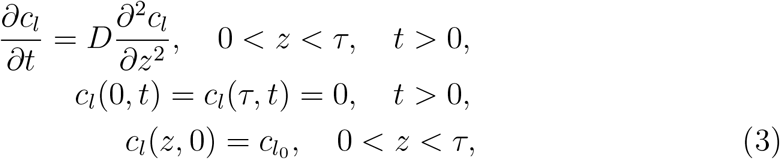

where *D* is the diffusivity of the therapeutic in the contact lens. The analytical solution of Eq. (3) was found by the method of separation of variables and is given by

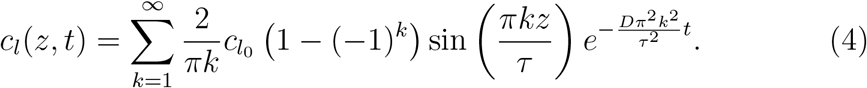

The cumulative release of the therapeutic from the contact lens was characterized by

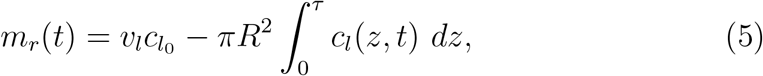

where *v*_l_ = *π R*^2^*τ* and *c*_*l*_(*z, t*) is given by Eq. (4). Note that *m*_r_(*t*) depends nonlinearly on the diffusion coefficient *D* via the exponential term in Eq. (4).

### 2.3. Parameter Estimation Algorithm

A two-step algorithm was implemented to estimate the diffusion coefficient of the therapeutic from a contact lens from experimental cumulative therapeutic release time series data.

#### Inputs

The inputs to the parameter estimate algorithm were

i. an experimental time series dataset of the cumulative release of the therapeutic from a contact lens placed in a vial of fluid, 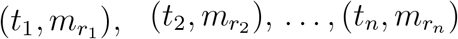;
ii. the initial amount of therapeutic loaded into the contact lens, 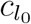;
iii. the radius, *R*, and the thickness, *τ*, of the contact lens (see Figure 1**A**).

When provided the total lens therapeutic absorption, like in Torres-Luna et al. (2020), the reported value was used to compute the initial condition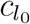; otherwise, 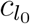 was approximated using the average of the last three time points of the therapeutic cumulative release data, i.e.,

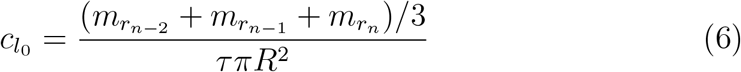

All contact lens radii, *R*, and thicknesses, *τ*, are reported in Table 2. All the values of initial mass of therapeutics in the lens 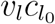 used are reported in Appendix C Table C.9.

**Step 1**. An approximation of the diffusion coefficient *D*, denoted by *D*_*exp*_, was found by fitting the experimental data to an exponential function, approximation of Eq. (5). More specifically, the time series cumulative release data was fitted to an exponential function given by

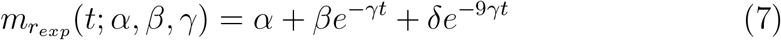

found by taking the first three terms of the series in Eq. (4) and plugging it into Eq. (5). The nonlinear least squares solver `lsqnonlin` in MATLAB 2022b was used to find values of *α, β*, and *γ*, that minimized the sum of the squares of the residuals given by

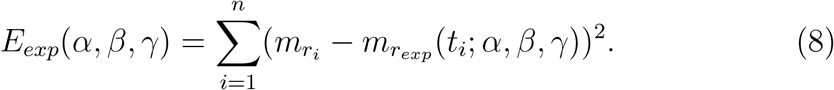

The approximation of diffusion coefficient *D*_*exp*_ was found by setting

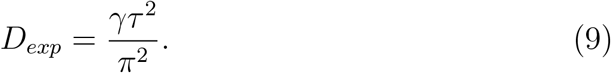

**Step 2**. An estimate *D*_*PDE*_ was found by minimizing a different nonlinear least squares error given by

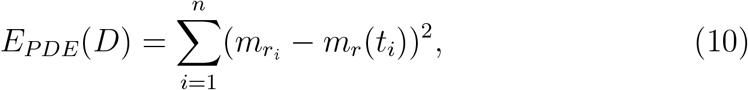

where *m*_*r*_(*t*) was approximated using the numerical solution of the partial differential equation problem in Eq. (3) and evaluating the integral of Eq. (5). To find the numerical solution of Eq. (3), the method of lines was used. The spatial partial derivatives were approximated by second order centered finite differences (Δ*z* = *τ* /*N*_*z*_, where *N*_*z*_ = 30) and the temporal partial derivative by a forward finite difference (Δ*t* < Δ*z*^2^/*D*). The integral in Eq. (5) was approximated using the trapezoidal rule.

Starting from the value of *D*_*exp*_, an interval of D containing *D*_*exp*_ was identified, see Figure 2**B**, and the minimum of the error *E*_*PDE*_ was found inside such interval. The algorithm was run until a minimum of Eq. (10) was reached and the corresponding diffusion coefficient was denoted as *D*_*PDE*_.

**Figure 2:**
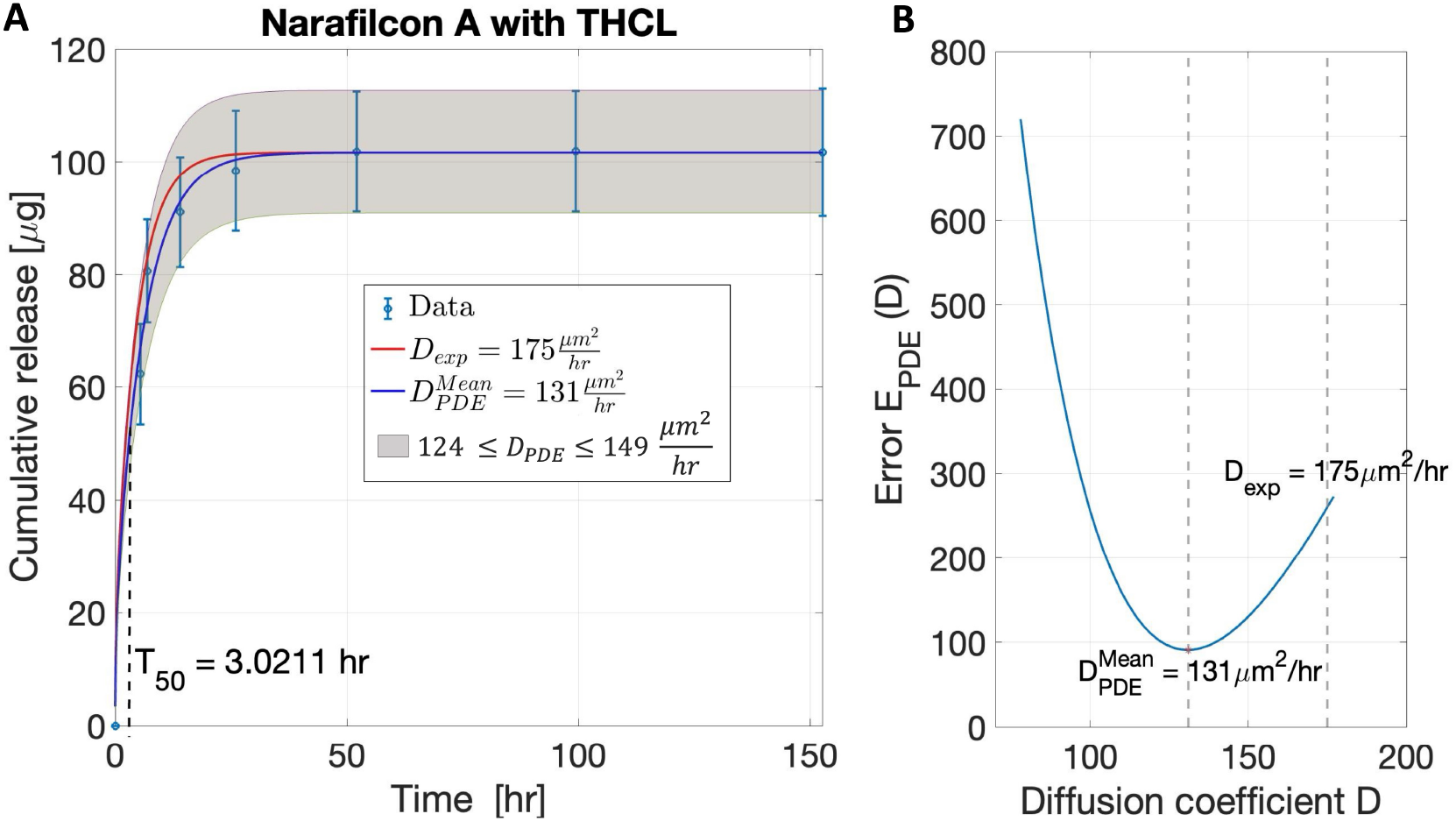
Parameter estimation for the release of tetracaine hydrochloride (THCL) from a narafilcon A contact lens. (**A**) The predicted cumulative therapeutic release *m*_*r*_ with *D* = *D*_*exp*_ (red curve) and 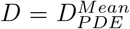 (blue curve). The shaded region is the predicted region of cumulative therapeutic release for the estimated range of diffusion coefficients. Data are presented as mean plus and minus the standard deviation. The dashed vertical line corresponds to the time that it takes for 50% of the loaded therapeutic to be released from the lens (*T*_50_). (**B**) The error, *E*_*PDE*_, as a function of *D*.

#### Outputs

Given the variability of the data, for each lens and therapeutic combination, a range of diffusion coefficients was estimated and a region (range) of cumulative therapeutic release was predicted. For each data set, the algorithm was performed to fit the average data of cumulative release and obtain the average value of *D*_*PDE*_, denoted as 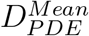. The average data plus and minus their standard deviation were fitted to obtain upper and lower bounds for *D*_*PDE*_.

The time at which the concentration of the therapeutic in the contact lens is equal to 50% of the initial amount loaded in the contact lens 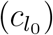 was found and was denoted as *T*_50_. For each lens and therapeutic combination, the MATLAB function `find` was used to search when *m*_*r*_(*t*) is within two decimal places of 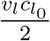. The value of *T*_50_ estimates the time that it takes for 50% of the loaded therapeutic to be released from the lens. The diffusivity, *D*, and time, *T*_50_, depend on each other; in general, the larger the value of the diffusion coefficient *D*, the faster the therapeutic will be released from the lens, i.e, the lower the value of *T*_50_.

Figure 2**A** shows the algorithm predicted cumulative therapeutic release of tetracaine hydrochloride (THCL) from a narafilcon A contact lens from Step 1 (red curve) and Step 2 (blue curve), and the experimental data (blue open circles) from Torres-Luna et al. (2020). The range of predicted *D*_*PDE*_ is depicted by the gray region in Figure 2**A**. Figure 2**B** shows the error between the numerical solution and the data, *E*_*PDE*_. The minimum of *E*_*PDE*_ occurred at 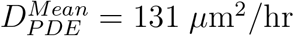, where the root mean square error, 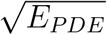, was less than 10% of the total amount of therapeutic released. The vertical dashed line in Figure 2**A** represents the predicted value of *T*_50_ for the release of THCL from a narafilcon A contact lens.

### 2.4. Statistical Analysis

A total of eight continuous variables were considered: the diffusion coefficient 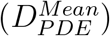 (found by fitting to the mean cumulative release data); the time to release 50% of the therapeutic from the contact lens (*T*_50_); the water content, thickness (*τ*), radius (*R*) of the contact lens; and the molecular volume, molecular mass, and density of the therapeutic.

Quantitative continuous variables were described by sample means, medians and standard derivations (SD). The normality of the sample data was first assessed using the Shapiro-Wilk’s test using the MATLAB function `swft`, with *p*-value > 0.05 and skewness between +1 and − 1 BenSaïda (2025). Mean comparisons of the normally distributed data sampled independently from two unrelated groups were carried out with a t-test using the MATLAB function `ttest2`. First, a two-tailed t-test was performed to assess the differences in means between the groups, and when the test was statistically significant, such differences were invested further using the upper- or lower-tailed t-test. In general, a *p*-value of < 0.05 was consider statistically significant. For non-normally distributed data sampled from two unrelated groups, a Mood’s median test was carried out to assess the difference in their respective median using the MATLAB function `mediatest` (Keine, 2025).

Since some of the variables were not normally distributed, the Kendall rank correlation coefficient (*r*_*Kendall*_) was used to evaluate the correlation between continuous variables. The MATLAB function `corrplot` was used to calculate correlations.

Multiple linear regression analysis was performed to create a predictive tool of 50% therapeutic release times for a combination of lens and therapeutic properties. The MATLAB function `regress` was used. The normality of the residuals was verified using the Shapiro-Wilk’s test, with *p*-value > 0.05 and skewness between +1 and -1. Data, whose corresponding residual did not follow a normal distribution, were considered outliers and were removed. An iterative approach was used to remove such outliers. A normal probability plot was created, using MATLAB function `normplot`, to compare the distribution of the residuals to the normal distribution. The data point furthest from normal probability plot was removed. The multiple linear regression analysis was performed again and the normality of the residuals were tested again using the Shapiro-Wilk’s test. If failed, the process was repeated again. Otherwise, the value of the coefficient of determination, denoted by *R*^2^, the corresponding *p*-value, the value of the regression coefficients, and the corresponding 90% confidence intervals (CI) were reported.

## 3. Results

Validation of the parameter estimation algorithm is presented first in Section 3.1, followed by the meta-analyses of the estimated diffusion coefficients in Section 3.2. Then a predictive tool is introduced to estimate release times of therapeutics from contact lenses in Section 3.3.

### 3.1. Validation of the Model and Algorithm

The algorithm was validated by comparing the estimated diffusion coefficients to those reported in the literature. Lanier et al. (2021) developed a mathematical model to estimate the partition coefficient (*K*) and diffusion coefficient (*D*) of a contact lens during *in vitro* uptake and release of the hydrophobic therapeutic cyclosporine. Diffusional transport of the therapeutic was assumed. The parameter 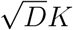 was fit to the *in vitro* soaking data, and the parameter *K* to the *in vitro* release data. *In vitro* experiments were conducted on five different lens types and then parameters were estimated for each lens type. The results of the parameter estimation algorithm herein were compared to the HEMA-based lenses (one of the five different types) as the HEMA lenses cumulative release profiles were close to sink conditions (as discussed in Lanier et al. (2021) and assumed here).

Table 4 reports the estimated diffusion coefficient, *D*_*PDE*_, and the estimated lens dimensions, and the initial loaded therapeutic mass used to calculate it. The assumed initial therapeutic mass loaded into the contact lens was estimated from the reported experimental loaded data (see Figure 4 of Lanier et al. (2021)). Specifically, the difference between the first reported drug mass in the vial to the last reported mass is used; such difference was assumed to be absorbed by the soaking lens. The estimated diffusion coefficient falls into the range of the Lanier et al. (2021)’s estimate. The error relative to the reported mean is 0.7%.

**Table 4:** Results of model validation. The input variables used to derive the diffusion coefficient estimated are reported and the estimate D_PDE_ is compared to the value of D reported in the literature: *a* = Lanier et al. (2021) and *b* = Pimenta et al. (2016).

| Lens<br>Drug | Thickness<br>$\tau$ [mm] | Radius<br>$R$ [mm] | Initial mass<br>loaded<br>$\pi R^2 \tau c_{l_0}$ [$\mu\text{g}$] | Estimated<br>value<br>$D_{PDE}$ [$\frac{\mu\text{m}^2}{\text{hr}}$] | Reported<br>value<br>$D$ [$\frac{\mu\text{m}^2}{\text{hr}}$] |
| --- | --- | --- | --- | --- | --- |
| HEMA<br>Cyclosporine | 0.15 | 6.0 | 20.00 | 15.80 | 15.91<br>±<br>1.90<sup>a</sup> |
| HEMA<br>Levofloxacin | 0.30 | 6.0 | 24.52 | 2780.00 | 2700.00<sup>b</sup> |

Pimenta et al. (2016) developed a mathematical model to estimate the diffusion coefficient in single-layered and multi-layered systems. The release of levofloxacin and chlorhexidine from HEMA-based discs was considered. The model included the two parameters *D* and *α*, which represent the diffusivity and the mass transfer between layers in the multi-layered lens system, respectively (Pimenta et al., 2016). In Pimenta et al. (2016), parameters *D* and *α* were estimated by comparing the numerical solution of the mathematical model to the experimental solution and then adjusting them accordingly. In validating the model presented herein in conparison to Pimenta et al. (2016), were considered only the single-layered lenses and the soluble drug levofloxacin. Chlorhexidine was not considered since it is insoluble.

Table 4 compares the predicted diffusion coefficient with their reported value of diffusivity for the combination of a single-layered HEMA contact lens with levofloxacin (see Table 1 in Pimenta et al. (2016)). It was assumed an initial mass of 24.52 mg levofloxacin was absorbed by the lens during the soaking process (same amount of levofloxacin released, sink boundary conditions). The estimated diffusion coefficient *D*_*PDE*_ = 2780 *µ*m^2^/hr was close to the diffusivity *D* = 2700 *µ*m^2^/hr reported by Pimenta et al. (2016); the relative error is 3%.

### 3.2. Meta-analysis of Estimated Diffusion Coefficients

Figure 3 shows the predicted ranges of *D*_*PDE*_ for all the combinations of contact lenses and therapeutics considered (see Table 1). The therapeutics, in the horizontal axis, are presented in order of ascending molecular mass (see Table 3). For each therapeutic, the lenses are divided in CH (orange names and dashed orange boxes) and SH (bold blue names and solid line blue boxes) and, within each lens type, listed in order of ascending water content (see Table 2). The lighter colored bar represents the portion of the range of *D*_*PDE*_ below 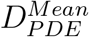 and the darker colored bar the portion of the range above. All the predicted values of 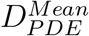 and the range of *D*_*PDE*_ are reported in Appendix C Table C.9.

**Figure 3:**
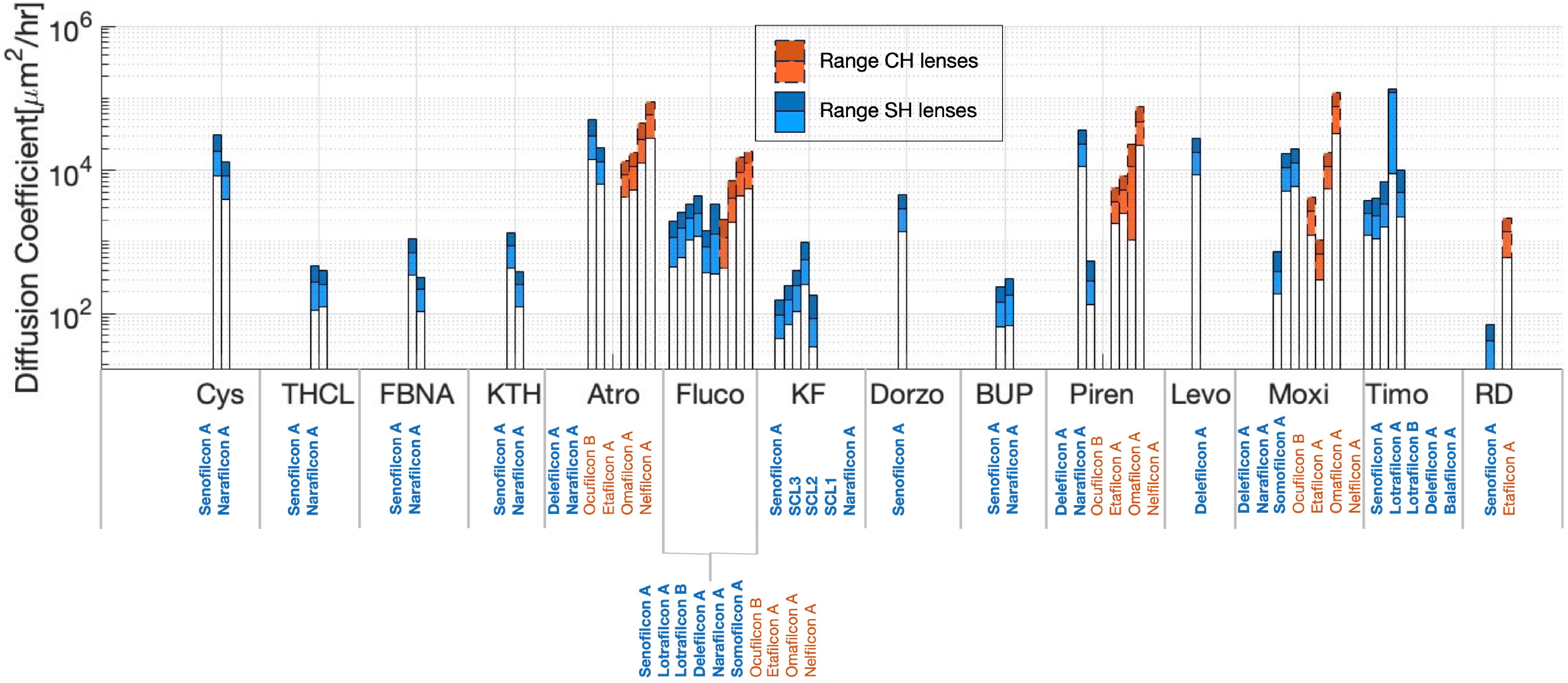
Predicted ranges of *D*_*PDE*_ for all the combinations of contact lenses and therapeutics considered. The lenses were divided into commercial hydrogel (CH, orange names and dashed orange boxes) and silicone hydrogel (SH, bold blue names and solid line blue boxes). The therapeutics considered were cysteamine (cys), tetracaine hydrochloride (THCL), flurbriprofen sodium (FBNA), ketorolac thromethamine (KTH), atropine (Atro), fluconazole (Fluco), ketotifen fumarate (KF), dorzolamide (Dorzo) bupivacaine(BUP), pirenzepine (Piren), levofloxacin (Levo), moxifloxacin (Moxi), timolol (Timo) and red dye (RD).

The predicted values of 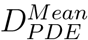, in logarithmic scale, are shown versus the contact lens and the therapeutic properties in Figure 4. SH lenses are reported in blue and CH lenses in solid orange, different symbols are used to identify the different FDA Groups. Correlations were computed between the 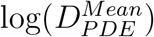, and the lens and therapeutic properties considered in Tables 2 and 3. Qualitatively, the predicted values of 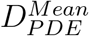, in logarithmic scale, was found to increase with water content (Figure 4**C**), and increase with therapeutic density (Figure 4**E**). Kendall’s rank correlation analysis shows that 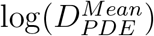 was positively correlated with water content (*r*_*Kendall*_ = 0.34, *p* < 0.001) and density of the therapeutic (*r*_*Kendall*_ = 0.23, *p* = 0.020). No statistically significant correlation was found with the remaining lens and therapeutic features.

**Figure 4:**
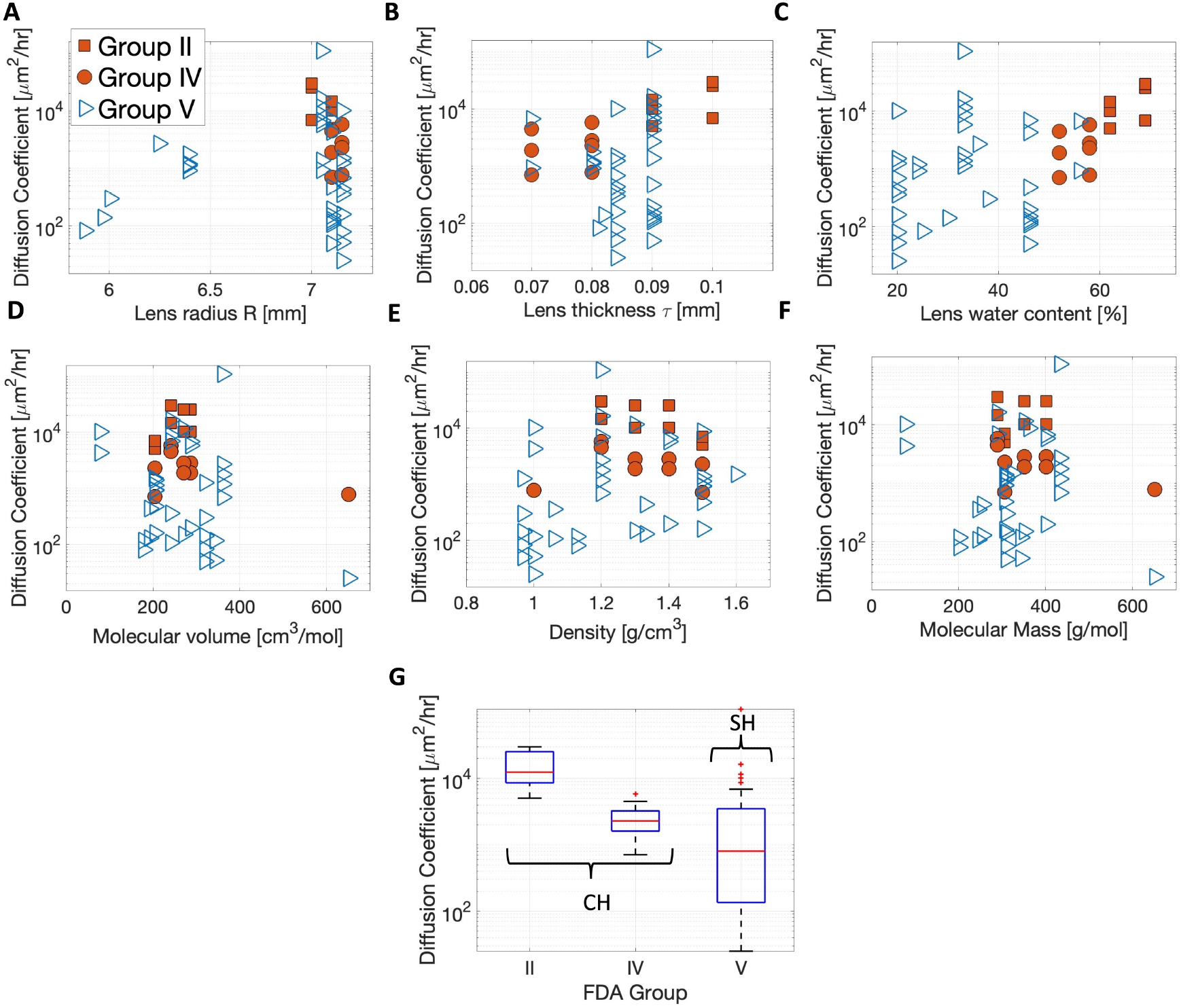
(**A**-**F**) The predicted values of 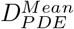 are shown on a logarithmic scale versus contact lens and therapeutic properties, and (**G**) the boxplot of 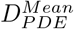 (shown on logarithmic scale) divided by FDA Groups. In **A**-**F**, silicon hydrogel (SH) lenses are reported in blue and commercial hydrogel (CH) lenses in orange. Different marker style symbols are used to represent the different FDA Groups.

FDA Groups were defined based on differences in water content (Groups I and III are low water content, Groups I and IV are high water content) and ionic charge (Groups I and III are nonionic, Groups III and IV are ionic). Figure 4**G** shows the boxplot of the predicted values of 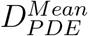 divided by FDA Groups. The sample mean, median, and standard deviation of 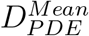 are reported in Table 5 for the CH lenses, SH lenses, and for the corresponding FDA Groups. The differences in 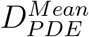 were investigated statistically between the type of lens, CH vs SH, and FDA Groups. Groups II (*N* = 8, Shapiro-Wilk’s test, *p* = 0.204) and IV (*N* = 9, Shapiro-Wilk’s test, *p* = 0.333) samples were normally distributed so t-test analysis results are reported in Table 5. CH lenses (*N* = 17, Shapiro-Wilk’s test, *p* = 0.001) and SH lenses (*N* = 36, Shapiro-Wilk’s test, *p* < 0.001) (same as Group V) samples were not normally distributed so Mood’s median analysis results are reported in Table 5. SH lenses median estimated diffusion coefficient was statistically different from the CH lenses median (*p* < 0.001). Group II, when compared Groups IV, showed a statistically higher mean 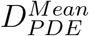 (*p* = 0.008). The median estimated diffusion coefficient for Groups IV and V were statistically different (*p* = 0.008).

**Table 5:** Mean, standard deviation (SD), and median of the estimated diffusion coefficient for the SH and CH lenses considered, and the corresponding FDA Groups. For normally distributed datasets (denoted by ^†^), *p*-values of the *t*-test for differences in means (≠, two-tailed) are reported, and when statistically significant (*p* < 0.05), the *p*-values of the upper or lower-tailed t-test are also reported. Mood’s median test, with *p*-value, are reported for non-normally distributed datasets. The symbol * denotes statistically significant *p*-values < 0.05.

| Estimated diffusion coefficient<br>$D_{PDE}^{Mean}$ | Silicone hydrogel (SH) lenses<br>(N=36) | Conventional hydrogel (CH) lenses<br>(N=17) | |
| --- | --- | --- | --- |
| Mean<br>± SD $\left[ \frac{\mu\text{m}^2}{\text{hr}} \right]$ | $5.656 \times 10^3$<br>$\pm 1.875 \times 10^4$ | $8.880 \times 10^3$<br>$\pm 9.384 \times 10^3$ | |
| Median $\left[ \frac{\mu\text{m}^2}{\text{hr}} \right]$ | $9.200 \times 10^2$ | $5.040 \times 10^3$ | |
| | same as **Group V** | **Group II<sup>†</sup>**<br>(N=8)<br>$1.593 \times 10^4$<br>$\pm 9.530 \times 10^4$<br>$1.231 \times 10^4$ | **Group IV<sup>†</sup>**<br>(N=9)<br>$2.613 \times 10^3$<br>$\pm 1.660 \times 10^3$<br>$2.289 \times 10^3$ |
| $t$-test, $p$-value | Group II<sup>†</sup> ≠ Group IV<sup>†</sup>, $p < 0.001^*$<br>(Group II<sup>†</sup> $>$ Group IV<sup>†</sup>, $p < 0.001^*$) | | |
| Mood's median test, $p$-value | SH ≠ CH, $p < 0.001^*$<br>Group II<sup>†</sup> ≠ Group V, $p = 0.008^*$<br>Group IV<sup>†</sup> ≠ Group V, $p = 0.008^*$ | | |

The differences in estimated diffusion coefficient 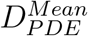 were studied for ionic lenses (*N* = 10), non-ionic lenses (*N* = 38), and unknown ionicity lenses (*N* = 5) and reported in Table 6. Mood’s median test was used as the non-ionic lens sample (Shapiro-Wilk’s test, *p* < 0.001) and unknown ionicities lens sample (Shapiro-Wilk’s test, *p* = 0.002) were not normally distributed. A statistically significant difference was found when comparing the median 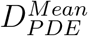 of ionic lenses with non-ionic lenses loaded with both positively and negatively changed therapeutics (*p* = 0.033). When comparing ionic, non-ionic, and unknown ionicity contact lenses loaded with only the positively charged therapeutics (see Table 3), no statistically significant differences were found in median of 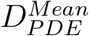 for the different lens ionicity groups. Not enough data were available to perform the same analysis when lenses were loaded only with negatively charged therapeutics (see Table 3).

**Table 6:** Mean, standard deviation (SD), and median of the estimated diffusion coefficients, 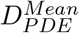, for the ionic, non-ionic, and unknown ionicities lenses. Mood’s median test for differences in medians (≠) are reported with *p*-value. The symbol * denotes statistically significant *p*-values < 0.05. The symbol ^†^ denotes the normally distributed dataset.

| **Estimated diffusion coefficient**<br>$D_{PDE}^{Mean}$ | **Ionic lenses<sup>†</sup>**<br>(N=10) | **Non-ionic lenses**<br>(N=38) | **Unknown ionicities lenses**<br>(N=5) |
| --- | --- | --- | --- |
| Mean | $2.622 \times 10^3$ | $8.277 \times 10^3$ | $1.632 \times 10^3$ |
| ±<br>SD $\left[\frac{\mu\text{m}^2}{\text{hr}}\right]$ | ±<br>$1.565 \times 10^3$ | ±<br>$1.880 \times 10^4$ | ±<br>$2.881 \times 10^3$ |
| Median $\left[\frac{\mu\text{m}^2}{\text{hr}}\right]$ | $2.494 \times 10^3$ | $1.319 \times 10^3$ | $3.000 \times 10^2$ |
| Mood's median test, $p$-value | Ionic<sup>†</sup> ≠ Non-ionic, $p = 0.033^*$<br>Ionic<sup>†</sup> ≠ Unknown ionicity, $p = 0.143$<br>Non-ionic ≠ Unknown ionicity, $p = 0.170$ | | |

### 3.3. Predictions of Therapeutic Release Times

A predictive tool was constructed for the logarithm of the mean 50% release time, log(*T*_50_), given lens and therapeutic properties, using a multiple linear regression analysis.

Correlation analysis was used to decide which lens and therapeutics properties should be considered in the multiple linear regression. The log(*T*_50_) dataset, containing all lens-therapeutic combinations, is reported in Appendix C Table C.9. Kendall’s rank correlation analysis shows that log(*T*_50_) was statistically significantly correlated with water content (*r*_*Kendall*_ = −0.43, *p* < 0.001) and therapeutic density (*r*_*Kendall*_ = −0.36, *p* = 0.003). Therefore, the explanatory variables in the regression used were the lens water content *w*_*c*_, where *w*_*c*_ ∈ [0, 100], and the therapeutic density *t*_*d*_, where the units of *t*_*d*_ are g/cm^3^.

The multiple linear regression model used is given by

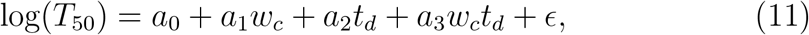

where *a*_*j*_, *j* = 0, .., 3 denote the regression coefficients and *ϵ* the error term. When fitted to all lens-therapeutic combinations (*N* = 53), the model’s coefficient of determination was *R*^2^ = 0.43; i.e., only approximately 43% of the variance in log(*T*_50_) is explained by lens water content and therapeutic density.

The three-dimensional plot of *T*_50_ (logarithmic scale) vs lens water content and therapeutic molecular volume is shown in Figure 5**A**. The data are divided into two groups depending on the molecular volume: small molecular volume between 75 cm^3^/mol and 275 cm^3^/mol (solid green, *N* = 32) and large molecular volume between 275 cm^3^/mol and 650 cm^3^/mol (purple, *N* = 21). Squares represent CH lenses and triangles represent SH lenses. Note that the two molecular volume ranges were chosen such that each of them includes both CH and SH lenses. Figure 5**A** shows that log(*T*_50_) qualitatively increases with water content and that such a trend might depend on the molecular volume of the therapeutic (different slope of the trend for the solid green symbols vs purple symbols). Recall that molecular volume was computed dividing the therapeutics molecular mass by its density.

**Figure 5:**
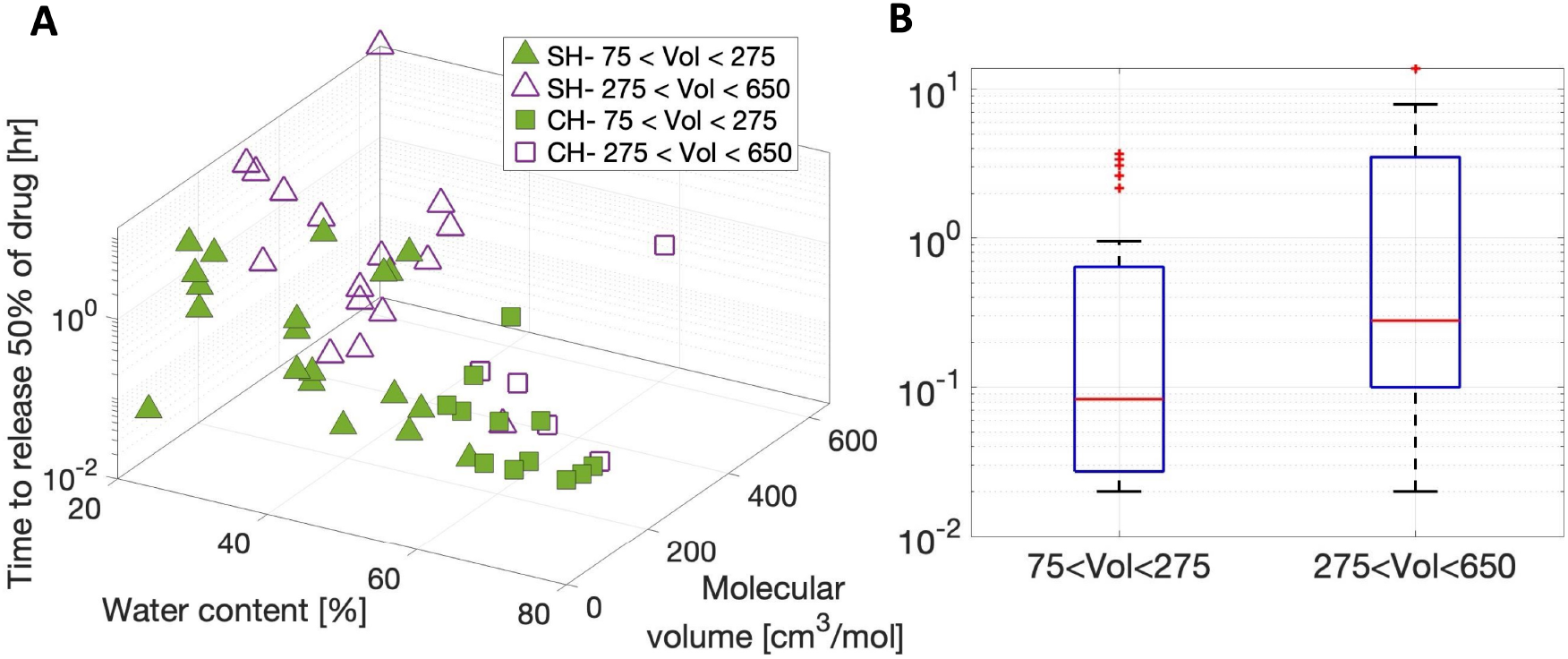
(**A**) Three-dimensional plot of *T*_50_ (logarithmic scale) vs lens water content and therapeutic molecular volume. The data are divided into two groups depending on the therapeutic molecular volume (Vol): volume between 75 cm^3^/mol and 275 cm^3^/mol (solid green) and volume between 275 cm^3^/mol and 650 cm^3^/mol (purple). Squares represent CH lenses and triangles represent SH lenses. (**B**) Comparison of the box plot of the *T*_50_ (logarithmic scale) for the therapeutic molecular volume groups considered.

Figure 5**B** compares the box plot of *T*_50_ (logarithmic scale) in the two groups of molecular volume considered. The Mood’s median test confirms the median log(*T*_50_) for the small molecular volume therapeutics was statistically different from the median log(*T*_50_) for large molecular volume therapeutics (*p* = 0.0378). For these reasons, the multiple linear regression analysis was then performed on the small molecular volume therapeutics and the large molecular volume therapeutics separately.

A graphical representation of the theoretical predictive tool (surface plot) for small molecular volume therapeutics (**A**) and for large molecular volume therapeutics (**B**) is shown in Figure 6. The outliers (or data excluded to ensure normality of residuals), the coefficient of determination, *R*^2^, with *p*-value, and the values of the regression coefficients, *a*_*j*_, and their 90% confidence intervals (CI_*j*_) are reported in Table 7. The coefficient of determination, *R*^2^, increased from 0.43, when considering all therapeutics together, to 0.64 when considering only the small molecular volume therapeutics and to 0.84 when considering only the large molecular volume therapeutics.

**Table 7:** Multiple linear regression model parameters for the two therapeutics molecular volume (Vol) groups, where *N* is the size of the dataset after removing the outliers. The outliers are lens-therapeutic combinations removed to ensure normality of residuals. Abbreviations are as follows: THCL: tetracaine hydrochloride, FBNA: flurbriprofen sodium, KTH: ketorolac thromethamine, Piren: pirenzepine, and Moxi: moxifloxacin. The coefficient of determination, with *p*-value, and the regression coefficients *a*_*j*_, with 90% confidence intervals (CI_*j*_), are reported. The symbol * denotes statistically significant *p*-values < 0.05.

|  | **75<Vol<275**<br>[cm<sup>3</sup>/mol]<br>(N=26) | **275<Vol<650**<br>[cm<sup>3</sup>/mol]<br>(N=19) |
| --- | --- | --- |
| Outliers:<br>Lens, Therapeutic | Narafilcon A, KTH<br>Etafilcon A, Piren<br>Elefilcon A, Piren<br>Narafilcon A, Piren<br>Narafilcon A, THCL<br>Narafilcon A, FBNA | Delefilcon A, Moxi<br>Narafilcon A, Moxi |
| Coefficient of<br>determination, $R^2$,<br>$p$-value | $R^2 = 0.64$, $p < 0.001^*$ | $R^2 = 0.84$, $p < 0.001^*$ |
| Regression<br>coefficient, $a_j$,<br>90% Confidence<br>interval ($CI_j$) | $a_0 = -0.5378$,<br>$CI_0 = [-7.2270, 6.1514]$<br>$a_1 = -0.0413$,<br>$CI_1 = [-0.2055, 0.1229]$<br>$a_2 = 0.5720$,<br>$CI_2 = [-4.4674, 5.6114]$<br>$a_3 = -0.0175$,<br>$CI_3 = [-0.141, 0.1061]$ | $a_0 = 18.2000$,<br>$CI_0 = [8.7262, 27.6738]$<br>$a_1 = -0.2003$,<br>$CI_1 = [-0.4162, 0.0155]$<br>$a_2 = -15.8068$,<br>$CI_2 = [-24.1743, -7.4393]$<br>$a_3 = 0.1527$,<br>$CI_3 = [-0.0270, 0.3325]$ |

**Figure 6:**
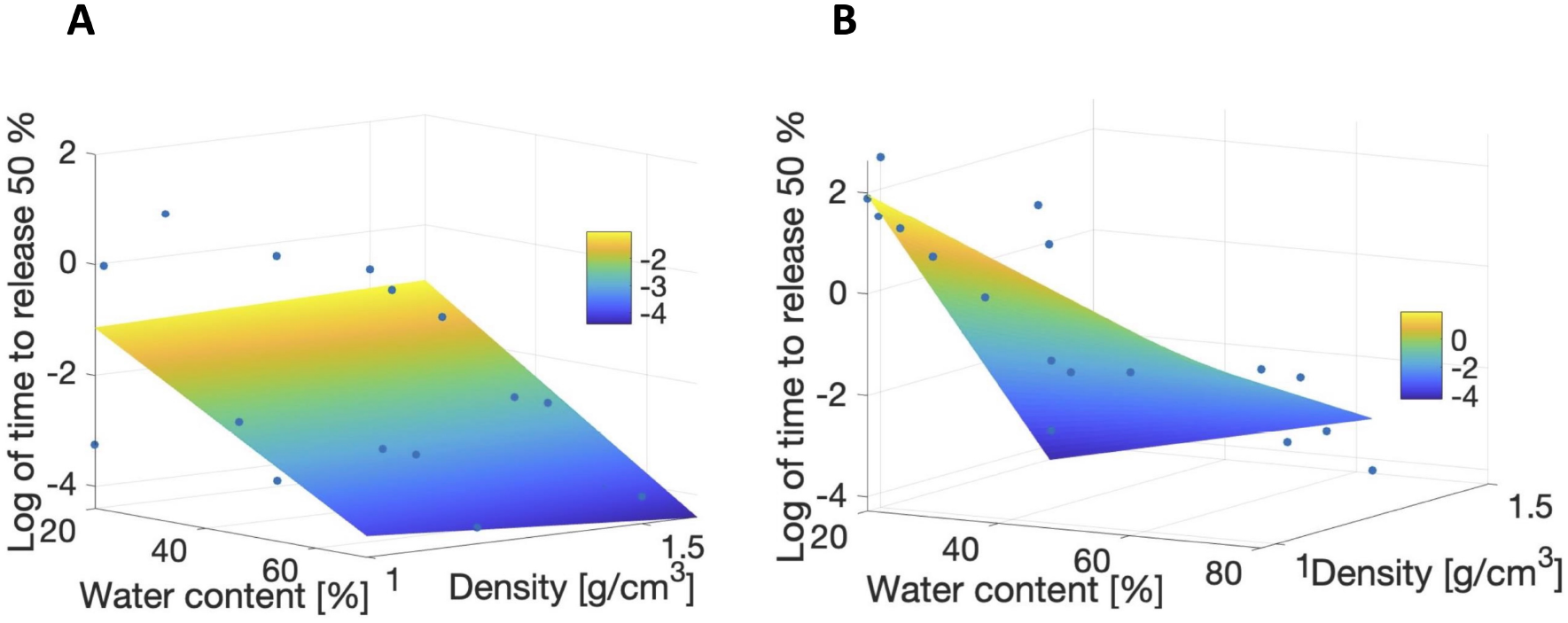
Graph of the predictive model for log(*T*_50_) (colored surface) vs data (blue circles) for (**A**) small-molecular-volume therapeutics and (**B**) large-molecular-volume therapeutics.

Figure 7 shows the predicted 50% release time for the small molecular volume therapeutic tetracaine hydrochloride (THCL, green line) and the large molecular volume therapeutic bupivacaine (BUP, purple line) for varying contact lens water content. The predicted time for 50% release is computed using Eq. (11) with either small molecular volume regression coefficients given in Table 7 and *t*_*d*_ = 1.13 g/cm^3^ (THCL, Vol= 178.00 cm^3^/mol) or large molecular volume regression coefficients and *t*_d_ = 0.99 g/cm^3^ (BUP, Vol= 343.07 cm^3^/mol). In general, the smaller molecular volume therapeutic was found to be released more quickly from contact lenses than the larger molecular volume therapeutic. As the water content of contact lens increases, the release time decreased.

**Figure 7:**
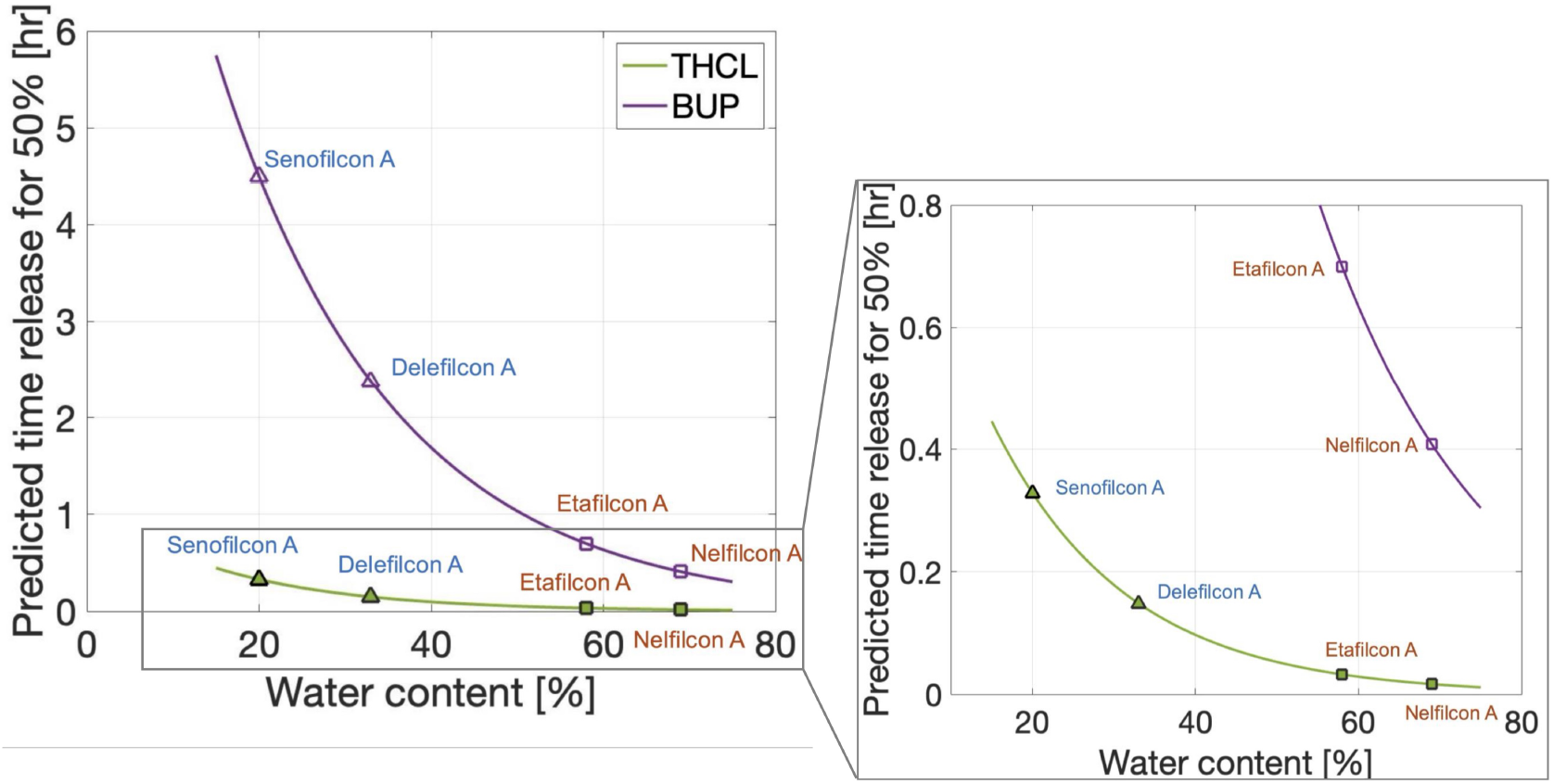
Predictive time of 50% release (*T*_50_) of THCL (small molecular volume therapeutic, green line) and BUP (large molecular volume therapeutic, purple line) varying contact lenses water content. Specific SH lenses senofilcon A and delefilcon A are denoted by triangle markers and blue labels, and CH lenses etafilcon A and nelfilcon A are denoted by square markers with orange labels.

Similarly, Figure 8 shows the predictive tools applied to the SH contact lens narafilcon A and the CH contact lens etafilcon A for different therapeutic densities. Figure 8**A** shows the dependence of *T*_50_ on therapeutic density for the narafilcon A lens (*w*_*c*_ = 46, blue line) and etafilcon A lens (*w*_*c*_ = 58, orange line) when considering small molecular volume therapeutics. The specific therapeutics plotted are tetracaine hydrochloride (THCL), flurbiprofen sodium (FBNA), ketrolomac tromethamine (KTH), atropine (Atro), fluconazole (Fluco), dorzolamide (Dorzo), levofloxacin (Levo) and pirenzepine (piren). Figure 8**B** displays the predicted 50% release time of large molecular volume therapeutics for SH contact lens narafilcon A (*w*_*c*_ = 46, blue line) and CH contact lens etafilcon A (*w*_*c*_ = 58, orange line). Specific therapeutics shown are ketotifen fumarate (KF), bupivacaine (BUP), moxifloxacin (Moxi) and timolol (Timo). Therapeutics, regardless of molecular volume, were released more quickly from a SH contact lens than from a CH contact lens. Small molecular volume therapeutics were released more quickly than large molecular volume therapeutics. As the density of the therapeutic increases, the therapeutic was released more quickly from a contact lens.

**Figure 8:**
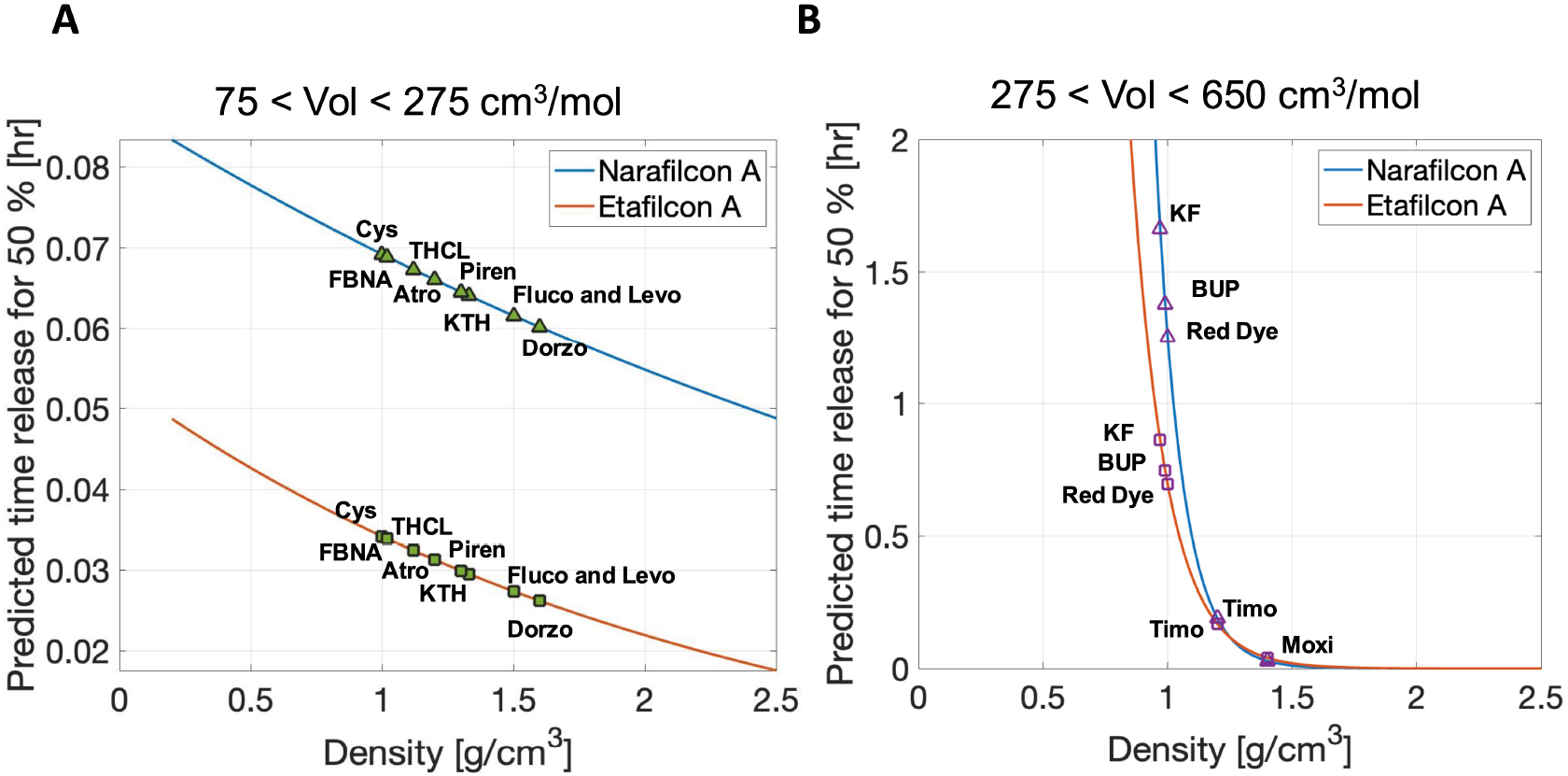
(**A**) Predictive 50% release time of small molecular volume therapeutics from SH contact lens narafilcon A (blue line) and CH contact lens etafilcon A (orange line) for varying therapeutic densities. Specific small molecular volume therapeutics shown are tetracaine hydrochloride (THCL), flurbiprofen sodium (FBNA), ketrolomac tromethamine (KTH), atropine (Atro), fluconazole (Fluco), dorzolamide (Dorzo), levofloxacin (Levo) and pirenzepine (piren). (**B**) Predictive 50% release time of large molecular volume therapeutics from SH contact lens narafilcon A (blue line) and CH contact lens etafilcon A (orange line) for varying therapeutic densities. Specific large molecular volume therapeutics shown are ketotifen fumarate (KF), bupivacaine (BUP), moxifloxacin (Moxi), and timolol (Timo).

## 4. Discussion

The meta-analysis conducted, leveraging the mathematical framework developed herein (see Figure 1), found that the kinetic release of therapeutics from contact lenses depends on both contact lens properties and therapeutics properties. In particular, the timing of the release was statistically significantly different depending on contact lens material (CH vs SH), contact lens ionicity, and therapeutic molecular volume (i.e., molecular mass divided by the density), and it was correlated with the contact lens water content and therapeutic molecular density.

In agreement with previous works, the lens material is important in characterizing release kinetics (Karlgard et al., 2003; Hui et al., 2017). In this work, therapeutics were found to be released faster, on average, from CH lenses than SH lenses. Table 5 reports that the median diffusivity of the CH lenses was statistically significantly different from the median diffusivity of the SH lenses, and Figure 5**A** shows CH lenses (squares) have shorter 50% release times than SH lenses (triangles). The timing of the therapeutic release was found to be one important piece of information when quantifying the dosage delivered to the ocular surface from a contact lens. Previous studies have reported that less therapeutics were released from a SH lens than a CH lens (Karlgard et al., 2003; Hui et al., 2017). Consequently, therapeutics could be released more slowly from a SH lens, but the amount of the therapeutic released could also be small.

Figure 3 shows the predicted diffusivity associated with the delefilcon A lens appears to behave differently from other SH lenses for certain drugs. The delefilcon A lens had predicted diffusivities very different from other SH lenses when loaded with pirenzepine and moxifloxacin, but like narafilcon A when loaded with atropine. The delefilcon A lens is marketed as a “hybrid” material (a hydrogel outer shell with a silicone core (Hui et al., 2017; Fan et al., 2020)). Thus, the likelihood of the release behavior of the hybrid lens mimicking other SH lenses depended on the therapeutic loaded into the lens.

The amount of water contained in the lens was found to correlate with therapeutic release timing. Specifically, high water CH content lenses (Group II) were found to have statistically significant greater mean diffusivity (smaller mean 50% release times) when compared to lower water SH content lenses (Group V). In general, water content was found to be an important predictor of the therapeutic release time regardless if the lens is a CH lens or SH lens. Figure 5**A** shows the smaller predicted 50% release times for the lenses with high water content. A similar trend was found among the three experimental silicone-based lenses fabricated by Xu et al. (2011) to study the release of ketotifen fumarate. The lens with the shortest release time of ketotifen fumarate had the highest measured equilibrium swelling ratio (a measure of the amount of water adsorbed by the contact lens). Additionally, the total amount of therapeutic loaded in and then released from a contact lens had also been found to depend on the water content of the lens (Karlgard et al., 2003; Hui et al., 2017).

Thang et al. (2023) argued that different levels of the composition and structures of hydrogel affect their swelling behavior which in turn influence the overall kinetics time release of therapeutics. Chatterjee et al. (2020) determined different compositions of the contact lenses can affect the release of certain therapeutics from contact lenses. In general, SH lenses with TRIS monomers exhibited lower cross-linking which on average lead to a higher swelling capacity (Chatterjee et al., 2020). Therefore, the release of therapeutics from TRIS containing SH lenses had an initial burst release. In the datasets studied, balafilcon A, lotrafilcon A, lotrafilcon B, SCL1, SCL2, and SCL3 were the only SH lenses with TRIS monomers (see Table 2). For all combination therapeutics, the results showed that most SH lenses with TRIS monomers had a larger diffusion coefficient when compared to other SH lenses without TRIS monomers (excluding the hybrid delefilcon A) such as narafilcon A and senofilcon A (see Figure 3).

Xu et al. (2011) reported the different compositions of the monomers in the lenses SCL1, SCL2, and SCL3 experimental contact lenses with their corresponding different swelling ratio. SCL3 had the smallest diffusion coefficient and the lowest equilibrium swelling ratio, while SCL2 had the largest diffusion coefficient and the highest equilibrium swelling ratio. Thus, an increase in the diffusion coefficient and, consequently faster therapeutic release, was predicted with an increase in swelling ratio which is consistent with the results provided by Xu et al. (2011).

Among the high-water content CH lenses (Groups II and IV), the non-ionic lenses (Group II) had a smaller median diffusivity than ionic lenses (Group IV) (see Table 6). Mood’s median test found the differences in the medians to be statistically significant. Consequently, the model predicted a slower median release of therapeutics from the non-ionic, high water content CH lenses than ionic. Prior works have found that surface charged or ionic CH contact lenses affected the amount of therapeutics released; specifically, ionic lenses released more therapeutics (Hui et al., 2017). For these reasons, could be hypothesized that for some therapeutics ionic, high water content CH lenses would release more therapeutics and do so more quickly than non-ionic, high water content CH lenses.

Etafilcon A, ocufilcon B and balafilcon A were the only ionic contact lenses (Groups IV and V) studied. In Figure 3, etafilcon A and ocufilcon B have the smallest diffusion coefficients among the subgroup of CH lenses when combined with positively charged drugs such as atropine, pirenzepine, and moxifloxacin. The negatively charged contact lenses might form electrostatic bonds with the cationic drugs causing a slower release of the drug which was captured in this work by smaller diffusion coefficients (Li and Mooney, 2016). Balafilcon A is the only ionic SH lens investigated. Saez-Martinez et al. (2021) investigated the microstructure of SH contact lenses and found balafilcon A to be the most ion permeable followed by lotrafilcon A and narafilcon A. Peng and Chauhan (2012) also argued that ion permeability could influence the transport of ionic drugs in SH contact lenses such that lenses with higher ionic permeability would have larger diffusivity. In Figure 3, a similar trend is observed for combination contact lenses with timolol released, where balafilcon A had a larger diffusion coefficient indicating faster release and could be hypothesized due to its large ionic permeability.

In this work, statistical differences were found in the median diffusivity of therapeutics considered in the ionic lenses (*N* = 10) and non-ionic lenses (*N* = 38), but no statistical significance in the median diffusivity of the therapeutics was observed when comparing either the ionic or non-ionic lenses with unknown ionicity contact lenses (*N* = 5). Soluri et al. (2012) found that contact lenses, such as etafilcon A, etafilcon B and balafilcon A, which are generally ionic contact lenses, had a larger uptake and release of therapeutic when compared to the non-ionic contact lenses, which supports the differences observed in the median diffusivities of ionic and non-ionic lenses.

The partition coefficient *K* of each lens-therapeutic combination were estimated using Eq. (1) and their values are reported in Table A.8. In this study, the combination of delefilcon A with atropine had the smallest partition co-efficient, while narafilcon A with ketotifen fumerate had largest partition coefficient. In comparison, in Figure 3, delefilcon A with atropine has one of the largest diffusion coefficient (fastest release of atropine) compared to narafilcon A with ketotifen which had one of the smallest diffusion coefficient (slowest release of ketotifen). On average, CH lenses had smaller partition coefficients in comparison to SH lenses with the exception of delefilcon A (the “hybrid” lens).

From the experiments evaluated in this study, the physical properties of therapeutics, that is, molecular volume (molecular mass/density) and density, were found to be explanatory variables of diffusivity and therapeutic release time. For the larger molecular volume therapeutics, the statistical model predicting the 50% release time described in Eq. (11) had an *R*^2^ value of 0.84 (see Table 7), i.e., capturing 84% of the variability of the data. The *R*^2^ value was 0.64 for the small molecular volume therapeutics. Small molecular volume therapeutics were found to release faster than larger molecular volume therapeutics. Karlgard et al. (2003) reported similar trends finding the smaller therapeutics to rapidly release from the contact lenses. Similar results were found in Hui et al. (2017), when comparing the amounts of atropine and pirenzepine released to the amount of dexamethasone released from the same lenses.

There are limitations to the present work. Simplifying assumptions were used to create the mathematical model herein. Most notably, the zero therapeutic concentration in the surrounding vial fluid. Consequently, the dynamics of the vial fluid were ignored. Prior works have quantified therapeutic concentrations in both the lens and surrounding fluid and, in the appropriate limits, predicted diffusivities are similar (Li and Chauhan, 2006; Pimenta et al., 2016; Lanier et al., 2021). Future work will tailor the mathematical model to predict temporal therapeutic release kinetics when the surrounding fluid is dynamic like the tear film.

The predicted model of the 50% release time for a given therapeutic/lens combination, given in Eq. (11), can be used in the preliminary stages of contact lens therapeutics delivery development to estimate the release time. The duration of the therapeutic release can be estimated; however, the amount of therapeutic release from the lens cannot be estimated as it requires additional information on the amount of therapeutics loaded. Additionally, the predicted model cannot quantify therapeutic loss. The initial amount of therapeutic loaded into the lens was assumed to be equal to the measured amount of therapeutic released from the lens.

If more experimental data were available, in particular, if all individual experimental data would be available rather than reported mean and standard deviations, the current statistical model developed in Eq. (11) could be extended to give a more robust interval estimate of 50% release time for a given therapeutic/lens combination via resampling methods such as bootstrapping.

## 5. Conclusions

A mathematical and statistical framework has been proposed herein to conduct a meta-analysis of the release of therapeutics from pre-soaked contact lenses placed in a vial of fluid. The framework was based on modeling therapeutic release via diffusion with sink boundary conditions. Using experimental data, the diffusivity and corresponding time for 50% of the therapeutics to be released were extracted and analyzed with respect to lens and therapeutics properties.

The water content of the lens was found to be positively correlated with the estimated diffusivity for all lens and therapeutic combinations studies. Consequently, the release time was hypothesized to be longer if a therapeutic was released from a SH lens when compared to a CH lens. The dependence of therapeutic release given the therapeutic charge and lens ionicity was found to be more nuanced, depending on if only positively charged therapeutics or only high water content lenses were considered or not. Finally, it was found that therapeutic molecular volume, therapeutic density, and contact lens water content could explain at least 64% of the variability in predicted 50% release times of therapeutics. The predicted model of the 50% release time for a given lens-therapeutic combination, given in Eq. (11), can be used in the preliminary stages of contact lens drug delivery development to estimate the release time.

## Acknowledgments

The authors would like to thank Dr. David Ross and Claire Canner for the help in the initial stages of the project and for the useful discussions. The authors would like to acknowledge Dr. Magdalena Cieslak for providing the raw datasets for some of the experiments and Dr. Carol Marchetti for the advice on some of the statistical analysis.

## Appendix A Partition Coefficient Estimates

Table A.8 reports all the numerical values, organized alphabetically by therapeutic, used to estimate the partition coefficient derived in Eq. (1).

**Table A.8:** Estimated values of the partition coefficient, see Eq. (1), for all lens-therapeutic combinations considered. The values of the loading therapeutics concentration and vial volume were extracted from the cited literature: *a* = Hui et al. (2017); *b* = Torres-Luna et al. (2020); *c* = Dixon et al. (2018); *d* = Hsu et al. (2015); *e* = Peng et al. (2010); *f* = Phan et al. (2016); *g* = Bajgrowicz et al. (2015); *h* = Torres-Luna et al. (2019); *i* = Xu et al. (2011); *j* = Dixon and Chauhan (2017); and *k* = Phan et al. (2021).

| Therapeutic | Lens | Loading conc<br>$c_{v_{loading}}$<br>[mg/ml] | Vial volume<br>$v_v$<br>[ml] | Lens vol<br>$v_l = \pi R^2 \tau$<br>[ml] | Partition coeff<br>$K$ | $\frac{\text{Lens vol}}{\text{Vial vol}}$<br>$\frac{v_l}{v_v}$<br>[%] |
| --- | --- | --- | --- | --- | --- | --- |
| Atropine | Delefilcon A<sup>a</sup> | 10.00 | 4.0 | 0.014 | 0.0019 | 0.35 |
|  | Narafilcon A<sup>a</sup> | 10.000 | 4.0 | 0.014 | 0.0021 | 0.36 |
|  | Ocufilcon B<sup>a</sup> | 10.000 | 4.0 | 0.013 | 0.0076 | 0.32 |
|  | Etafilcon A<sup>a</sup> | 10.000 | 4.0 | 0.013 | 0.0080 | 0.32 |
|  | Omafilcon A<sup>a</sup> | 10.000 | 4.0 | 0.014 | 0.0052 | 0.36 |
|  | Nelfilcon A<sup>a</sup> | 10.000 | 4.0 | 0.015 | 0.0020 | 0.38 |
| Bupivacaine | Senofilcon A<sup>b</sup> | 1.000 | 10.0 | 0.013 | 13.0376 | 0.13 |
|  | Narafilcon A<sup>b</sup> | 1.000 | 10.0 | 0.014 | 19.5144 | 0.14 |
| Cysteamine | Senofilcon A<sup>c</sup> | 50.000 | 3.0 | 0.013 | 0.9192 | 0.45 |
|  | Narafilcon A<sup>c</sup> | 50.000 | 3.0 | 0.014 | 0.9996 | 0.47 |
| Dorzolamide | Senofilcon A<sup>d</sup> | 0.800 | 3.5 | 0.013 | 12.0636 | 0.39 |
| Fluconazole | Senofilcon A<sup>e</sup> | 0.700 | 2.0 | 0.013 | 6.9860 | 0.67 |
|  | Lotrafilcon A<sup>e</sup> | 0.700 | 2.0 | 0.012 | 6.9830 | 0.60 |
|  | Lotrafilcon B<sup>e</sup> | 0.700 | 2.0 | 0.012 | 8.9742 | 0.60 |
|  | Delefilcon A<sup>f</sup> | 1.000 | 4.8 | 0.014 | 10.5269 | 0.29 |
|  | Narafilcon A<sup>f</sup> | 1.000 | 4.8 | 0.014 | 5.6290 | 0.30 |
|  | Somofilcon A<sup>f</sup> | 1.000 | 4.8 | 0.013 | 6.3846 | 0.28 |
|  | Ocufilcon B<sup>f</sup> | 1.000 | 4.8 | 0.013 | 14.9514 | 0.26 |
|  | Etafilcon A<sup>f</sup> | 1.000 | 4.8 | 0.013 | 12.0924 | 0.27 |
|  | Omafilcon A<sup>f</sup> | 1.000 | 4.8 | 0.014 | 9.0005 | 0.30 |
|  | Nelfilcon A<sup>g</sup> | 1.000 | 4.8 | 0.015 | 3.0853 | 0.32 |
| Fluobriprofen sodium | Senofilcon A<sup>h</sup> | 0.200 | 3.0 | 0.013 | 49.1691 | 0.45 |
|  | Narafilcon A<sup>h</sup> | 0.200 | 3.0 | 0.014 | 45.5571 | 0.47 |
| Ketotifen fumarate | Senofilcon A<sup>a</sup> | 0.300 | 10.0 | 0.013 | 44.1014 | 0.13 |
|  | SCL3<sup>i</sup> | 1.945 | 5.0 | 0.009 | 18.6872 | 0.18 |
|  | SCL2<sup>i</sup> | 1.945 | 5.0 | 0.009 | 23.2798 | 0.18 |
|  | SCL1<sup>i</sup> | 1.945 | 5.0 | 0.010 | 27.2340 | 0.19 |
|  | Narafilcon A<sup>a</sup> | 0.300 | 10.0 | 0.014 | 44.1298 | 0.14 |
| Ketorolac | Senofilcon A<sup>a</sup> | 1.200 | 3.0 | 0.013 | 8.2319 | 0.45 |

Table A.8: Estimated values of the partition coefficient, see Eq.
| Therapeutic | Lens | Loading conc<br>$c_{v_{loading}}$<br>[mg/ml] | Vial volume<br>$v_v$<br>[ml] | Lens vol<br>$v_l = \pi R^2 \tau$<br>[ml] | Partition coeff<br>$K$ | Lens vol<br>Vial vol<br>$\frac{v_l}{v_v}$<br>[%] |
| --- | --- | --- | --- | --- | --- | --- |
| tromethamine | Narafilcon A<sup>a</sup> | 1.200 | 3.0 | 0.014 | 7.6045 | 0.48 |
| Levofloxacin | Delefilcon A<sup>i</sup> | 1.500 | 3.0 | 0.014 | 0.8624 | 0.47 |
| Moxifloxacin | Delefilcon A<sup>g</sup> | 1.000 | 4.8 | 0.014 | 1.6297 | 0.29 |
|  | Narafilcon A<sup>g</sup> | 1.000 | 4.8 | 0.014 | 2.1655 | 0.30 |
|  | Somofilcon A<sup>g</sup> | 1.000 | 4.8 | 0.013 | 2.6676 | 0.28 |
|  | Ocufilecon B<sup>g</sup> | 1.000 | 4.8 | 0.013 | 17.1895 | 0.26 |
|  | Etafilecon A<sup>g</sup> | 1.000 | 4.8 | 0.013 | 17.5985 | 0.27 |
|  | Omafilecon A<sup>g</sup> | 1.000 | 4.8 | 0.014 | 9.7811 | 0.30 |
|  | Nelfilcon A<sup>g</sup> | 1.000 | 4.8 | 0.015 | 2.5900 | 0.32 |
| Pirenzepine | Delefilcon A<sup>b</sup> | 1.000 | 4.0 | 0.014 | 0.0142 | 0.35 |
|  | Narafilcon A<sup>b</sup> | 1.000 | 4.0 | 0.014 | 0.0161 | 0.36 |
|  | Ocufilecon B<sup>b</sup> | 1.000 | 4.0 | 0.013 | 0.0253 | 0.32 |
|  | Etafilecon A<sup>b</sup> | 1.000 | 4.0 | 0.013 | 0.0265 | 0.32 |
|  | Omafilecon A<sup>b</sup> | 1.000 | 4.0 | 0.014 | 0.0372 | 0.36 |
|  | Nelfilcon A<sup>b</sup> | 1.000 | 4.0 | 0.015 | 0.0182 | 0.38 |
| Red dye | Senofilcon A<sup>k</sup> | 1004.800 | 2.0 | 0.013 | 1.6533 | 0.67 |
|  | Etafilecon A<sup>k</sup> | 1004.800 | 2.0 | 0.013 | 3.7044 | 0.64 |
| Tetracaine hydrochloride | Senofilcon A<sup>a</sup> | 0.200 | 10.0 | 0.013 | 41.4192 | 0.13 |
|  | Narafilcon A<sup>a</sup> | 0.200 | 10.0 | 0.014 | 35.6946 | 0.14 |
| Timolol | Senofilcon A<sup>e</sup> | 0.800 | 2.0 | 0.013 | 8.8011 | 0.67 |
|  | Lotrafilcon A<sup>e</sup> | 0.800 | 2.0 | 0.012 | 4.7859 | 0.60 |
|  | Lotrafilcon B<sup>e</sup> | 0.800 | 2.0 | 0.012 | 7.4657 | 0.60 |
|  | Delefilcon A<sup>j</sup> | 5.000 | 3.0 | 0.014 | 1.7667 | 0.47 |
|  | Balafilcon A<sup>e</sup> | 0.800 | 2.0 | 0.011 | 14.8461 | 0.55 |

## Appendix B Justification of Sink Boundary Conditions

The mass of the therapeutic in the contact lens at time *t, m*_*l*_(*t*), was given by

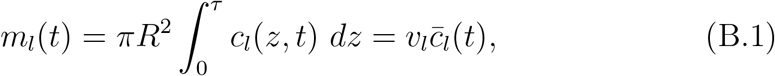

where *v*_*l*_ denotes the contact lens volume and 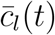 is the average therapeutic concentration in the contact lens. The contact lens volumes were estimated by assuming each contact lens is a cylinder with radius given by *R*, and height given by *τ*, both reported in Table 2, i.e. *v*_*l*_ = *πR*^2^*τ*. Therefore, the change of the contact lens therapeutic mass in time is computed as 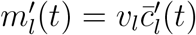.

The mass of the therapeutic in the vial at time *t* was given by

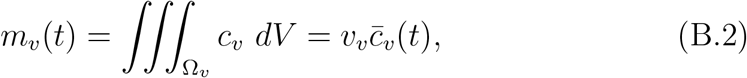

where Ω_*v*_ denotes the domain of the vial solution, *v*_*v*_ the volume of the solution in the vial and 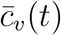 is the average therapeutic concentration in the vial. Therefore, the rate of change of the vial therapeutic mass in time is computed as 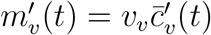.

Since the therapeutic mass was conserved in the experimental system and initially, *c*_*v*_|_*t*=0_ = 0, the therapeutic mass in the vial *m*_*v*_(*t*) = *m*_*l*_(0) − *m*_*l*_(*t*). So 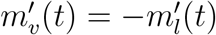.Thus, the rate of change of the average therapeutic vial concentration is given by

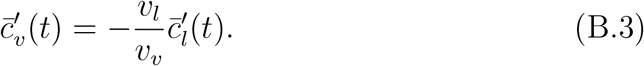

For all experiments analyzed, the ratio of the lens volume to the vial volume 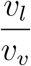 was less than 0.7% (see Table A.8 in Appendix A). Consequently, 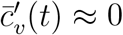, so the average therapeutic vial concentration was approximately constant during the experiment, and because the therapeutic vial concentration was initially zero, *c*_*v*_ ≈ 0.

Note that, using the notation introduced in this section, the cumulative release of the therapeutic from the contact lens defined in Eq. (5), can be expressed as *m*_*r*_(*t*) = *m*_*l*_(0) − *m*_*l*_(*t*).

## Appendix C Additional Results

Table C.9 reports the mean and range of the initial therapeutic mass loaded into the contact lens; the mean 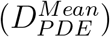 and range of the estimated diffusion coefficient, *D*_*PDE*_; and the predicted 50% release time of the therapeutic, *T*_50_, assuming the diffusion coefficient of the therapeutic is 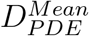.

**Table C.9:** The value of the initial therapeutic loaded mass, 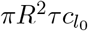, used to find the reported mean and range of the estimated diffusion coefficient, *D*_*PDE*_. The predicted 50% release time, T_50_, of therapeutic assuming the diffusivity of the therapeutic is given by the reported mean value, 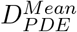. Literature referenced: *a* = Hui et al. (2017); *b* = Torres-Luna et al. (2020); *c* = Dixon et al. (2018); *d* = Hsu et al. (2015); *e* = Phan et al. (2016); *f* = Torres-Luna et al. (2019); *g* = Xu et al. (2011); *h* = Dixon and Chauhan (2017); *i* = Bajgrowicz et al. (2015); *j* = Phan et al. (2021); and *k* = Peng et al. (2010);.

| Therapeutic | Lens | Initial loaded therapeutic mass<br>$v_l c_{l_0} = \pi R^2 \tau c_{l_0}$<br>[$\mu\text{g}$] | Estimated diffusion coefficient<br>$D_{PDE}$<br>[$\frac{\mu\text{m}^2}{\text{hr}}$] | 50% release time<br>$T_{50}$<br>[hr] |
| --- | --- | --- | --- | --- |
| Atropine | Delefilcon A<sup>a</sup> | mean: 0.27<br>range:<br>0.25-0.28 | mean: 16300<br>range:<br>13990-20000 | 0.0210 |
|  | Narafilcon A<sup>a</sup> | mean: 0.36<br>range:<br>0.35-0.38 | mean: 6900<br>range:<br>6300-7500 | 0.0600 |
|  | Ocufilcon B<sup>a</sup> | mean: 0.96<br>range:<br>0.94-0.99 | mean: 4515<br>range:<br>4200-4800 | 0.0540 |
|  | Etafilcon A<sup>a</sup> | mean: 1.03<br>range:<br>0.10-1.05 | mean: 5808<br>range:<br>5400-6200 | 0.0522 |
|  | Omafilcon A<sup>a</sup> | mean: 0.74<br>range:<br>0.71-0.77 | mean: 14580<br>range:<br>12600-17300 | 0.0202 |
|  | Nelfilcon A<sup>a</sup> | mean: 0.31<br>range:<br>0.29-0.33 | mean: 29850<br>range:<br>27900-31700 | 0.0201 |
| Bupivacaine | Senofilcon A<sup>b</sup> | mean: 175.80<br>range:<br>170.33-181.27 | mean: 80<br>range:<br>65-92 | 4.3100 |
|  | Narafilcon A<sup>b</sup> | mean: 278<br>range:<br>232.59-320.96 | mean: 117<br>range:<br>67-122 | 3.3800 |
| Cysteamine | Senofilcon A<sup>c</sup> | mean: 619.7<br>range:<br>569.7-669.7 | mean: 10200<br>range:<br>8300-12700 | 0.0390 |
|  | Narafilcon A<sup>c</sup> | mean: 712.4<br>range:<br>684.0-740.8 | mean: 4305<br>range:<br>3880-4870 | 0.0923 |

| Therapeutic | Lens | Initial loaded<br>therapeutic<br>mass<br>$v_l c_{l_0} = \pi R^2 \tau c_{l_0}$<br>[$\mu\text{g}$] | Estimated<br>diffusion<br>coefficient<br>$D_{PDE}$<br>[$\frac{\mu\text{m}^2}{\text{hr}}$] | 50%<br>release<br>time<br>$T_{50}$<br>[hr] |
| --- | --- | --- | --- | --- |
| Dorzolamide | Senofilcon A<sup>d</sup> | mean: 122<br>range:<br>119.4-124.6 | mean: 1500<br>range:<br>1400-1600 | 0.2615 |
| Fluconazole | Senofilcon A<sup>e</sup> | mean: 65.94<br>range:<br>64.48-67.39 | mean: 690<br>range:<br>444-762 | 0.5000 |
|  | Lotrafilcon A<sup>e</sup> | mean: 58.60<br>range:<br>57.29 - 59.64 | mean: 920<br>range:<br>609-1122 | 0.3700 |
|  | Lotrafilcon B<sup>e</sup> | mean: 75.13<br>range:<br>73.96-76.28 | mean: 1130<br>range:<br>1055-1202 | 0.2700 |
|  | Delefilcon A<sup>e</sup> | mean: 147.86<br>range:<br>142.19-153.53 | mean: 1387<br>range:<br>1163-1813 | 0.0816 |
|  | Narafilcon A<sup>e</sup> | mean: 80.19<br>range:<br>77.99-82.38 | mean: 478<br>range:<br>367-606 | 0.0849 |
|  | Somofilcon A<sup>e</sup> | mean: 86.09<br>range:<br>81.15-91.03 | mean: 927<br>range:<br>357-2071 | 0.0219 |
|  | Ocufilcon B<sup>e</sup> | mean: 189.33<br>range:<br>174.40-204.26 | mean: 710<br>range:<br>430-936 | 0.0864 |
|  | Etafilcon A<sup>e</sup> | mean: 155.29<br>range:<br>125.58-184.99 | mean: 2288<br>range:<br>1830-3062 | 0.0212 |
|  | Omafilcon A<sup>e</sup> | mean: 128.22<br>range:<br>107.09-149.35 | mean: 5040<br>range:<br>4451-5587 | 0.0215 |
|  | Nelfilcon A<sup>e</sup> | mean: 47.47<br>range:<br>32.68-62.26 | mean: 6964<br>range:<br>5460-5587 | 0.0230 |
| Flubriprofen<br>sodium | Senofilcon A<sup>f</sup> | mean: 132.7<br>range:<br>126.5-138.9 | mean: 356<br>range:<br>337-385 | 0.9582 |
|  | Narafilcon A<sup>f</sup> | mean: 129.8<br>range:<br>124.6-135 | mean: 107.6<br>range:<br>107.1-108 | 3.6800 |
| Ketotifen<br>fumarate | Senofilcon A<sup>b</sup> | mean: 178.40<br>range:<br>151.56-193.37 | mean: 52<br>range:<br>45-61 | 6.6375 |
|  | SCL3<sup>g</sup> | mean: 319.62<br>range:<br>301.54-337.36 | mean: 83<br>range:<br>70-95 | 3.8600 |
|  | SCL2<sup>g</sup> | mean: 415.52<br>range:<br>387.81-443.23 | mean: 141<br>range:<br>108-150 | 2.3400 |
|  | SCL1<sup>g</sup> | mean: 503.81<br>range:<br>470.04-513.00 | mean: 300<br>range:<br>250-422 | 1.1500 |
|  | Narafilcon A<sup>b</sup> | mean: 188.60<br>range:<br>176.68-200.52 | mean: 50<br>range:<br>35-95 | 7.9000 |
| Ketrorolac<br>tromethamine | Senofilcon A<sup>f</sup> | mean: 133.2<br>range:<br>127.8-138.6 | mean: 436<br>range:<br>434-440 | 0.7859 |
|  | Narafilcon A<sup>f</sup> | mean: 130<br>range:<br>135-140 | mean: 129<br>range:<br>126-132 | 3.0800 |
| Levofloxacin | Delefilcon A<sup>h</sup> | mean: 4500<br>range:<br>4408-4592 | mean: 8755<br>range:<br>8700-10405 | 0.2615 |
| Moxifloxacin | Delefilcon A<sup>i</sup> | mean: 22.89<br>range:<br>20.15-25.64 | mean: 5792<br>range:<br>5190-5800 | 0.0703 |
|  | Narafilcon A<sup>i</sup> | mean: 30.85<br>range:<br>27.54-34.16 | mean: 196<br>range:<br>190-330 | 2.0337 |
|  | Somofilcon A<sup>i</sup> | mean: 35.97<br>range:<br>34.77-37.18 | mean: 6760<br>range:<br>6038-6800 | 0.0315 |
|  | Ocufilcon B<sup>i</sup> | mean: 217.67<br>range:<br>214.04-221.31 | mean: 1450<br>range:<br>1228-1550 | 0.1200 |
|  | Etafilcon A<sup>i</sup> | mean: 226<br>range:<br>223.18-226.50 | mean: 373<br>range:<br>300-401 | 0.1100 |
|  | Omafilcon A<sup>i</sup> | mean: 139.34<br>range:<br>136.91-141.76 | mean: 5880<br>range:<br>5540-5983 | 0.0400 |
|  | Nelfilcon A<sup>i</sup> | mean: 39.85<br>range:<br>37.69-42.00 | mean: 43000<br>range:<br>32555-44000 | 0.0200 |
| Pirenzepine | Delefilcon A<sup>a</sup> | mean: 0.20<br>range:<br>0.19-0.21 | mean: 11600<br>range:<br>11200-13300 | 0.0315 |
|  | Narafilcon A<sup>a</sup> | mean: 0.23<br>range:<br>0.22-0.24 | mean: 151<br>range:<br>135-251 | 2.6300 |
|  | Ocufilcon B<sup>a</sup> | mean: 0.32<br>range:<br>0.29-0.35 | mean: 1890<br>range:<br>1800-2100 | 0.1400 |
|  | Etafilcon A<sup>a</sup> | mean: 0.34<br>range:<br>0.30-0.38 | mean: 2820<br>range:<br>2520-3100 | 0.8400 |
|  | Omafilcon A<sup>a</sup> | mean: 0.53<br>range:<br>0.50-0.57 | mean: 10200<br>range:<br>1055-11500 | 0.0517 |

Table C.9: The value of the initial therapeutic loaded mass, $\pi R^2 \tau c_{l_0}$ , used to find the reported mean and range of the estimated diffusion coefficient, $D_{PDE}$ .
| Therapeutic | Lens | Initial loaded therapeutic mass<br>$v_l c_{l_0} = \pi R^2 \tau c_{l_0}$<br>[$\mu\text{g}$] | Estimated diffusion coefficient<br>$D_{PDE}$<br>[$\frac{\mu\text{m}^2}{\text{hr}}$] | 50% release time<br>$T_{50}$<br>[hr] |
| --- | --- | --- | --- | --- |
|  | Nelfilcon A<sup>a</sup> | mean: 0.28<br>range: 0.26-0.30 | mean: 25300<br>range: 22380-29300 | 0.0200 |
| Red dye | Senofilcon A<sup>j</sup> | mean: 22400<br>range: 20400-24400 | mean: 25<br>range: 17-30 | 13.8400 |
|  | Etafilcon A<sup>j</sup> | mean: 47800<br>range: 45100-50500 | mean: 776<br>range: 607-800 | 0.3100 |
| Tetracaine hydrochloride | Senofilcon A<sup>b</sup> | mean: 111.70<br>range: 105.37 - 116.08 | mean: 160<br>range: 110-200 | 2.1633 |
|  | Narafilcon A<sup>b</sup> | mean: 101.70<br>range: 90.91 -112.72 | mean: 131<br>range: 124-149 | 3.0211 |
| Timolol | Senofilcon A<sup>k</sup> | mean: 94.94<br>range: 74.95-114.93 | mean: 1250<br>range: 1240-1325 | 0.2820 |
|  | Lotrafilcon A<sup>k</sup> | mean: 45.79<br>range: 25.79-65.78 | mean: 1190<br>range: 1098-1826 | 0.2614 |
|  | Lotrafilcon B<sup>k</sup> | mean: 71.43<br>range: 51.44-91.42 | mean: 1760<br>range: 1609-3649 | 0.1764 |
|  | Balafilcon A<sup>k</sup> | mean: 131.11<br>range: 127.51-134.70 | mean: 2700<br>range: 2227-5316 | 0.1500 |

## Notes

### Competing Interest Statement

Dr. Phan is a cofounder of OcuBlink Inc., a Canadian startup that develops in vitro eye models for testing contact lenses.

